# A liquid biopsy platform for detecting gene-gene fusions as glioma diagnostic biomarkers and drug targets

**DOI:** 10.1101/2020.02.25.963975

**Authors:** Vikrant Palande, Rajesh Detroja, Alessandro Gorohovski, Rainer Glass, Charlotte Flueh, Marina Kurtz, Shira Perez, Dorith Raviv Shay, Tali Siegal, Milana Frenkel-Morgenstern

## Abstract

Gliomas account for about 80% of all malignant brain tumours. Diagnosis is achieved by radiographic imaging followed by tumour resection, to determine tumour cell type, grade and molecular characteristics. Glioblastoma multiforme (GBM) is the most common type of glioma, and is uniformly fatal. The median survival of treated GBM patients is 12-15 months. Standard modalities of therapy are unselective and include surgery, radiation therapy and chemotherapy, while precision medicine has yet to demonstrate improvements in disease outcome. We therefore selected GBM as a model to develop a precision medicine methodology for monitoring patients using blood plasma circulating cell-free DNA (cfDNA). Currently, tumour heterogeneity, clonal diversity and mutation acquisition are the major impedances for tailoring personalized therapy in gliomas in general, and particularly in GBM. Thus, a liquid biopsy diagnostics platform based on cfDNA sequencing may improve treatment outcome for GBM patients, by guiding therapy selection. In this study, we processed from 27 patients with glioma, 27 plasma samples for cfDNA isolation and 5 tissue biopsy samples for tumour DNA isolation. From a control group of 14 healthy individuals, 14 plasma samples were processed for cfDNA isolation. In glioma patients, cfDNA concentration was elevated compared to controls. Point mutations found in glioma tissue biopsies were also found in the cfDNA samples (95% identity). Finally, we identified novel chimeric genes (gene-gene fusions) in both tumour and cfDNA samples. These fusions are predicted to alter protein interaction networks, by removing tumour suppressors and adding oncoproteins. Indeed, several of these fusions are potential drug targets, particularly, NTRK or ROS1 fusions, specifically for crizotinib analogues (like entrectinib and larotrectinib) with enhanced penetration of the central nervous system. Taken together, our results demonstrate that novel druggable targets in gliomas can be identified by liquid biopsy using cfDNA in patient plasma. These results open new perspectives and abilities of precision medicine in GBM.

## Introduction

Gliomas are primary malignant brain tumours that account for about 30% of all brain tumours and 80% of malignant brain tumours^1–3^. Glioblastoma multiforme (GBM) is the most common type of glioma, and is uniformly fatal; the median survival of treated patients is approximately 12-15 months^4,5,6^. The gold standard method for detecting gliomas is radiographic evaluation by magnetic resonance imaging (MRI) scanning^3,7,8^ followed by tissue diagnosis attained either by biopsy or during surgery. Tissue sampling must be obtained to determine tumour cell type, histologic grade and molecular characteristics^7,9^. The MRI contrast-enhancing region of the tumour is the target for surgical resection as part of GBM treatment^10^, which includes combined chemo-radiation therapy. In about 40% of GBM, *O*^6^-methylguanine DNA methyltransferase (MGMT) promoter is methylated. This renders the cancer more susceptible to temozolomide, an alkylating agent that methylates DNA, and that constitutes a standard chemotherapy for GBM^11^. Current methods for tumour monitoring (e.g., MRI and CT) cannot provide real time actionable information for determining therapy response or for evolving the molecular landscape of the heterogeneous cancer cell population^12^. Furthermore, even a more precise diagnostic method such as molecular analysis of tumour biopsy may not entirely represent a heterogeneous tumour, or newly acquired mutations during the course of the disease^13,14^. A liquid biopsy platform that uses circulating cell-free DNA (cfDNA) may overcome the limitations posed by glioma tumour heterogeneity, and may provide a diagnostic means, and possibly even guide precision medicine for GBM^12,15,16^.

Liquid biopsy is a newly emerging non-invasive cancer diagnostic technique that potentially provides an alternative to surgical biopsies. Liquid biopsy provides various information about a tumour from simple blood, urine, saliva, serum or plasma samples^13,17–20^. This technique uses cells or cell components circulating in blood or urine, such as circulating cell-free DNA(cfDNA)^21,22^, cell-free RNA(cfRNA)^23–28^, extracellular vesicles^29–31^, circulating proteins^32,33^ and circulating tumour cells^34–40^. These components are continuously released from the tumour and healthy tissues into the bloodstream as a result of secretion, rapid apoptosis and necrosis^41,42^. These moieties can be screened for tumour specific markers that may be useful in cancer diagnosis, monitoring or prognosis.^17,19,43^

CfDNA constitutes free-floating small fragments of DNA in the blood plasma, which result from apoptotic cell death^44^. Although sparsely studied, remarkably elevated cfDNA has been documented in patients with solid tumours compared to those with non-neoplastic diseases^32,45–47^. Of all the cfDNA fragments present in a cancer patient’s plasma, 85% are 166-bp, 10% are 332-bp and 5% are 498-bp in length^21^ (Fig. 1). By contrast, larger fragments of cfDNA, ∼10,000 bp in length, in the blood of cancer patients, are most likely of necrotic origin^41,42,48–50^ (Fig. 1). Using next-generation sequencing analysis, *Snyder* et al.,^51^ identified some bias in the cfDNA fragmentation pattern, which was affected by nucleosome occupancy and transcription factor binding. The latter protects DNA from nuclease digestion during apoptosis, potentially providing a clue to cell type origin^51^. CfDNA has been found in patients with diverse types of neoplasms and metastatic disease^52,53^. This has led to the identification of specific genetic markers bearing varying degrees of specificity and sensitivity. However, these markers have yet to prove useful in diagnostics^54–61^. Deep sequencing of plasma DNA in patients with various cancers suggests that cfDNA is representative of the entire tumour genome, and can be an accurate reflection of tumour heterogeneity and acquired mutations^13,14,61–63^. As such, in breast, colon, ovarian and melanoma cancers, cfDNA levels have been found to be clinically useful, with an inverse relation between cfDNA levels and survival^55,64–68^. By contrast, a major problem arises in the analysis of cfDNA from brain tumours due to low cfDNA levels in plasma. The unique localization and anatomic features of the brain^54^, appear to reduce the frequency of detectable cfDNA by 60% in medulloblastoma, and by 90% in low-grade glioma, compared to other advanced systemic tumours. This is substantially lower than the levels detected in colon, breast, lung, prostate, renal and thyroid cancers^69–72^ Thus, detecting cfDNA in glioma samples for clinical implementation remains a complex challenge^54^.

**Fig. 1.**
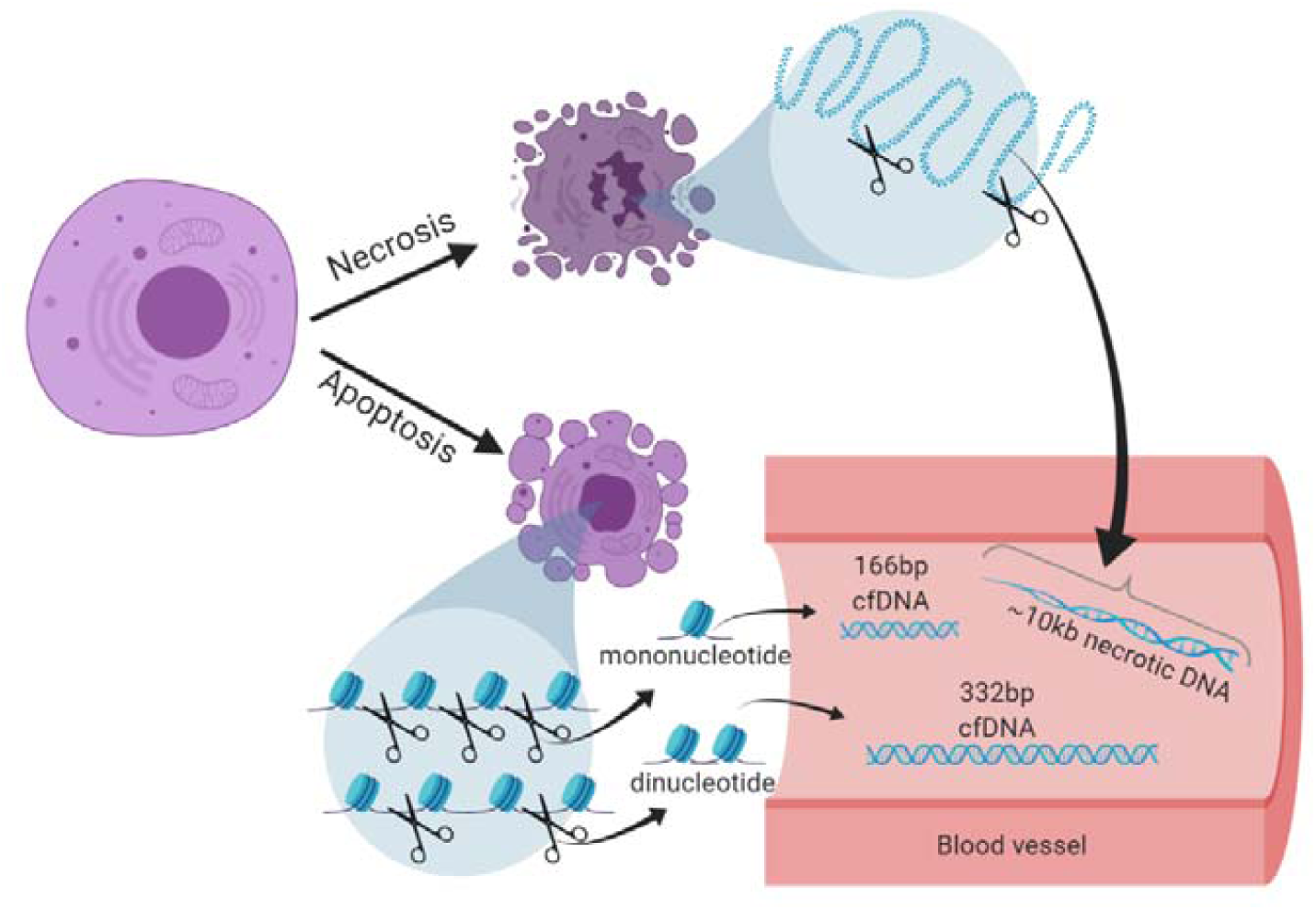
Different sizes of cfDNA found in the blood as a result of apoptosis and necrosis.

Precision medicine in glioma is in its infancy. Only a handful of mutations have been associated with disease type and prognosis, and no mutations are known to guide treatment modality. Thus, the World Health Organization presently classifies tumours of the central nervous system (CNS) based on their histological and molecular parameters, and on associations of mutations with disease prognosis and therapy^73,74^. However, these mutations have not been shown to guide precision medicine treatment of glioma. Diagnostic genes in brain tumours include isocitrate dehydrogenase (*IDH*), *H3K27M, RELA* fusion, wingless (WNT)-activated, sonic hedgehog (SHH)-activated and *C19MC*-altered^75–78^. A typical molecular signature that was extensively studied in oligodendroglial tumours is a chromosomal 1p/19q co-deletion^79–81^. The exclusive presence of the 1p/19q co-deletion in oligodendroglial tumours renders it a required biomarker for oligodendroglioma diagnosis, including hTERT promoter mutation^82–84^.

In regard to fusion genes as molecular markers, chromosomal aberrations play a crucial role in the initial steps of tumorigenesis^85–88^. This is especially true for translocations and their corresponding gene fusions^89–92^, as they disrupt cellular regulatory mechanisms. These fusion genes are highly tissue specific and can be used as effective biomarkers in cancer diagnosis^85–88,93,94^. For example, *TMPRSS2-ERG* fusion genes have been detected in 40-80% of prostate cancers^89–91^. Moreover, the *BCR-ABL* fusion gene is most commonly observed in CML.^92^ Overall, around 90% of lymphomas and nearly half of all forms of leukaemia harbour translocation-induced gene fusions.^95^ The *EML4-ALK* gene fusion plays a crucial role in the development of epithelial cancers and lung cancer^96^, while the *C11orf95-RELA* fusion is characteristic of ependymoma grade II and grade III tumours (World Health Organization classification)^73^. Thus, the presence of the *C11orf95-RELA* fusion gene serves as an important biomarker for glioma subtype diagnosis^87^. Additional fusion genes may also serve as novel molecular markers for cancer diagnosis, yet their detection in liquid biopsy has not been systematically demonstrated, particularly in brain tumours.

Current technologies such as next-generation sequencing and digital droplet PCR can be applied with high sensitivity towards rare mutation detection^97–103^, and can thus be used for identifying recurrent translocations and gene fusions in many cancer types including gliomas^104–107^. We have collected more than 40,000 unique fusion transcripts (of more than 40 cancer types) in our ChiTaRS-5.0 database of Chimeric Transcripts and RNA-Seq data^108,109^. This is the largest collection of chimeric transcripts (of cancer chromosomal translocations and RNA trans-splicing) known today, including sense-antisense transcripts (SaS chimeras)^109^. Moreover, we have collated data on 1,207 known druggable fusion genes from PubMed articles using our text-mining method, ProtFus^110^. In the present study we sequenced cfDNA of glioma patients; and assessed plasma concentration, mutation patterns and novel fusion genes. By comparing to our existing fusion database, we identified potential liquid biopsy biomarkers, and also drug targets and their corresponding druggable fusions. The findings may help guide precision medicine in glioma therapy.

## Results

### Elevated cfDNA concentration in the plasma of glioma patients

We hypothesized that cfDNA concentration might differ between individuals with gliomas and a non-cancer cohort. Thus, we collected 14 blood samples from healthy controls and 27 blood samples from patients with gliomas (25 glioblastoma, 2 low-grade glioma). The latter were matched to 27 brain biopsies of the same patients. For each patient, we extracted cfDNA from plasma, genomic DNA (gDNA) from white blood cells (WBC), and tumour DNA (tDNA) from brain biopsies. First, we assessed the cfDNA plasma concentration in the control cohort as ranging from 0 to 7.62ng per ml of plasma (Fig. 2). Next, we isolated cfDNA in 26 of 27 samples of glioma patients, and achieved a 96% sensitivity level. The Qubit concentrations of cfDNA in these 26 patients ranged between 12.6ng and 137ng per ml of plasma (Fig. 2). Thus, all the patients had concentrations that were higher than those of all of the control group (p-value <0.0001, t-test). Next, we examined the size of the cfDNA molecules in all the samples. The Bioanalyzer DNA High Sensitivity assay showed that in all 26 glioma patients, and also in all 14 healthy control samples, cfDNA had a major peak at, or close to, 166bp (accounting for 85% of the circulating cfDNA) and a smaller peak at, or close to, 332bp (accounting for 10% of the cfDNA; Fig. 3A). This concurs with the reference sizes of cfDNA fragments in plasma^21,111^. Moreover, the cfDNA sequencing library peak was measured at 291bp, which indicates successful ligation of a 125bp adapter to 166bp cfDNA (Fig. 3B). Thus, a liquid biopsy methodology can generate high quality results, enabling analysis of the cfDNA that is likely derived from apoptotic cells (rather than necrosis). Accordingly, overall plasma cfDNA levels appear to discriminate a tumour-bearing cohort (including patients with brain tumours) from a control cohort.

**Fig. 2.**
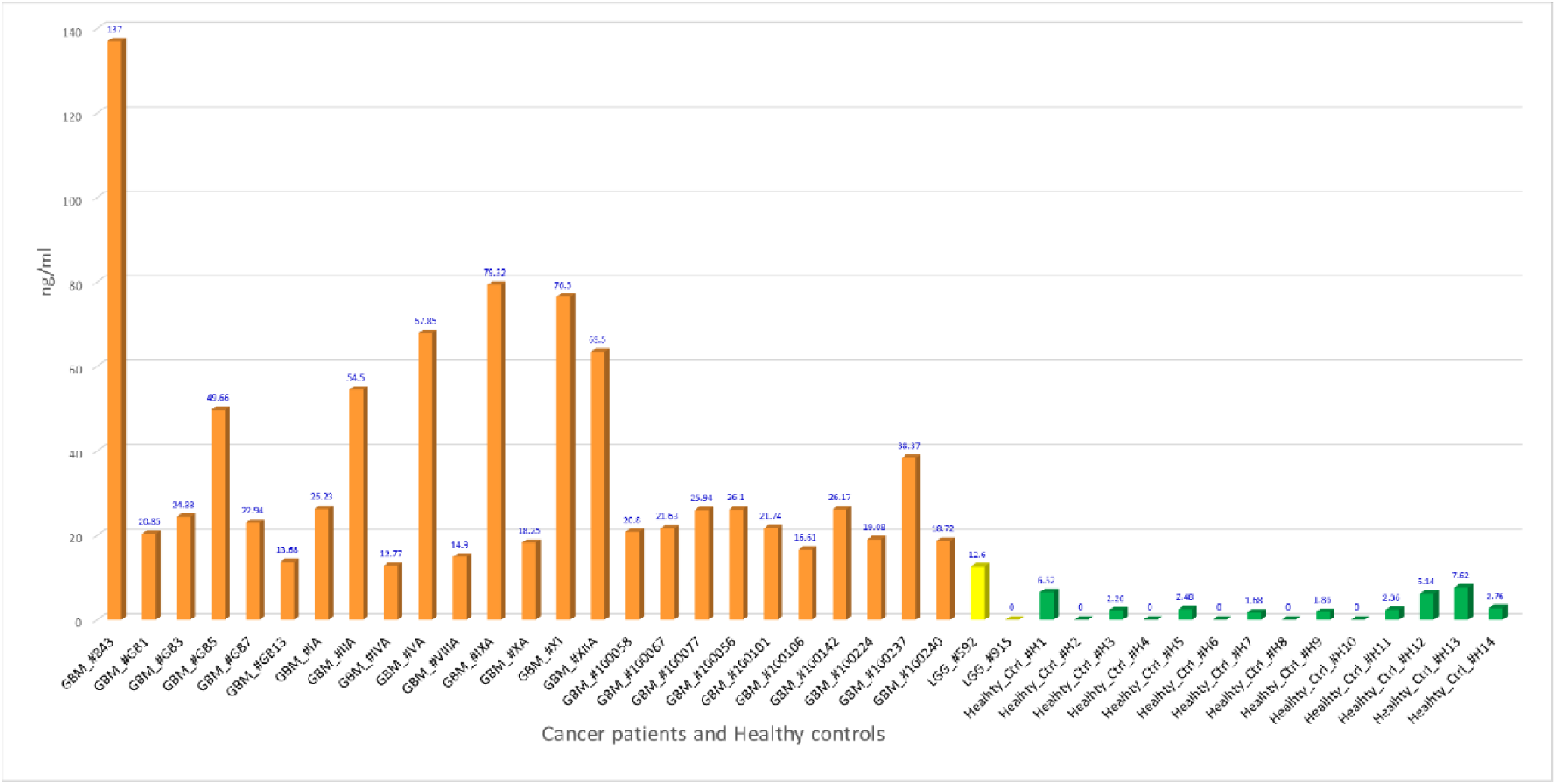
Cell-free DNA (cfDNA) concentrations in glioma cancer patients vs. healthy controls. The graph shows comparatively high cfDNA concentrations in 1 ml plasma of 25 glioblastoma patients (orange), 2 low-grade glioma patients (yellow) and 14 control samples (green).

**Fig. 3.**
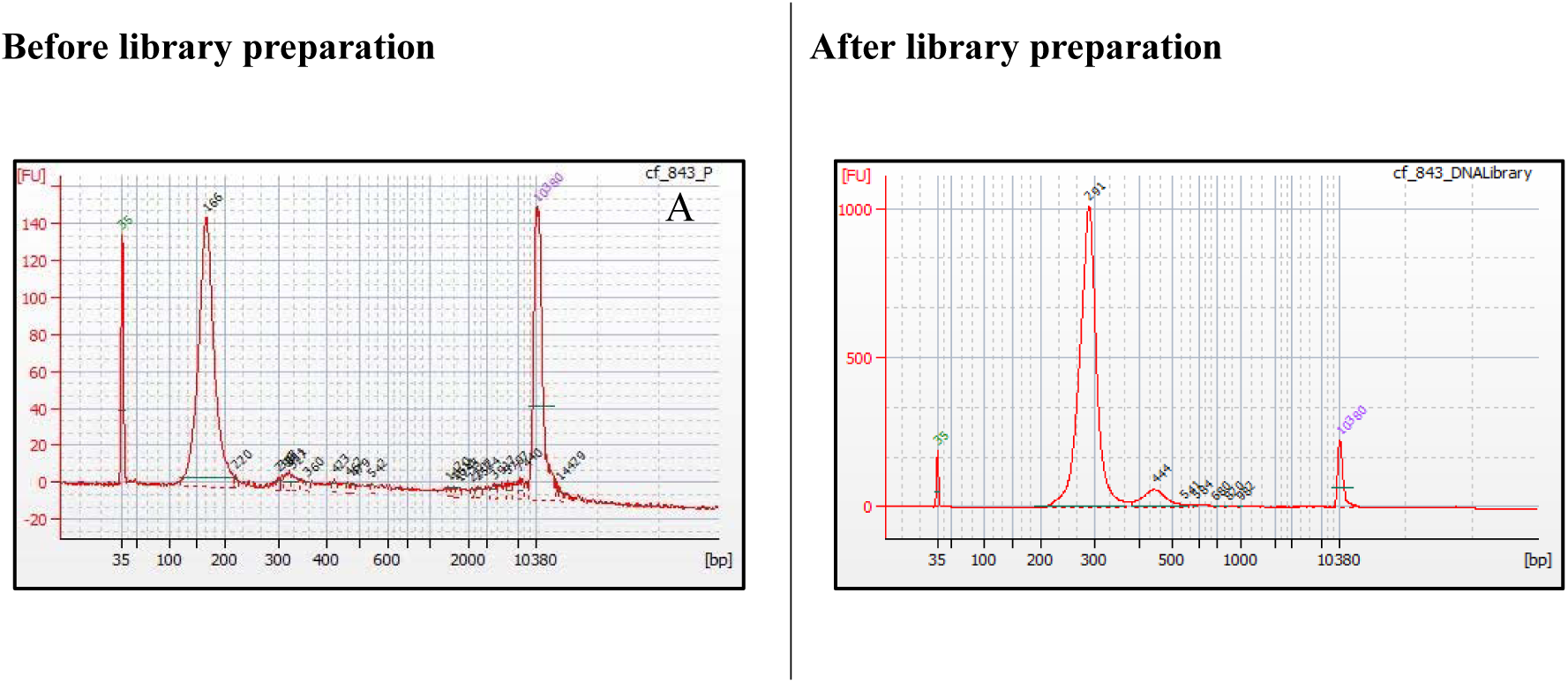
Bioanalyzer assay electropherograms of cell-free DNA (cfDNA) before and after the next-generation sequencing (NGS) library preparation step. **(A)** cfDNA isolated from glioblastoma patient #843, enriched at fragment size 166bp. **(B)** cfDNA NGS library enriched at the expected fragment size 291bp (corresponding to 166bp of cfDNA + 125bp NGS adapters), confirming its successful NGS library preparation.

**Fig. 4.**
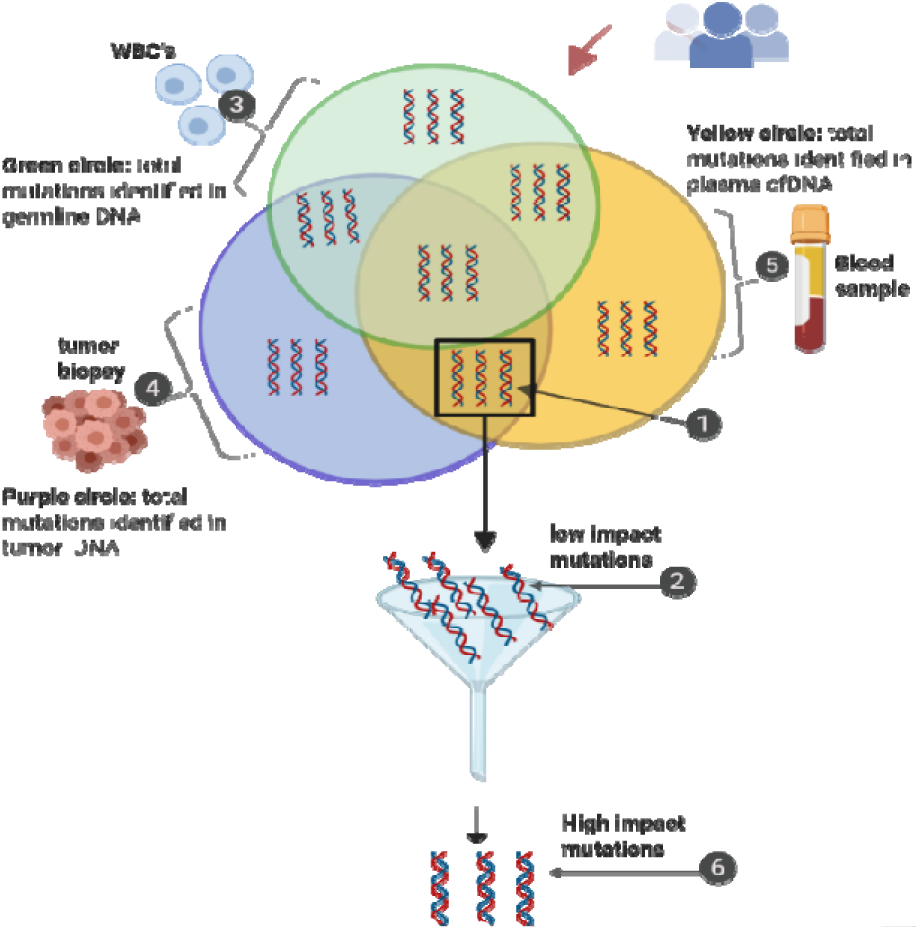
Variant analysis method used to identify high impact variants.

### Mutation analysis of glioma cfDNA data

To confirm that the elevated cfDNA in the plasma of patients with glioma is derived from cancer cells, we tested for the presence of mutations in both cfDNA and tDNA. We sequenced 23 cfDNA samples (14 glioma and 9 controls) using a whole genome sequencing procedure (see Materials and Methods) with 5x-10x coverage (at least 50 million paired-end (PE) reads per sample). In addition, for five glioma samples, we sequenced their respective tDNA (10x coverage, 60 million PE reads) and normal genomic DNA (from WBC) (∼60 million PE reads). We first removed the SNPs present in gDNA, then sorted mutations into “cfDNA only”, “tDNA only”, and “both cfDNA and gDNA”. We found for all five GBM patients, namely #GB1, #GB3, #GB5, #GB7 and #GB13, shared mutations between their cfDNA and tDNA, with 90% selectivity and 80% sensitivity (FDR<1%, Table 1). These results indicate that in GBM, cfDNA derives from the brain tumours (presumably, efflux from extracellular space into the plasma).

**Table 1.** The numbers of high impact alterations by each consequence type that were commonly identified in cell-free DNA (cfDNA) and tumour DNA (tDNA) of glioblastoma patients #GB1, #GB3, #GB5, #GB7 and #GB13

| Consequence type<br>(Sequence Ontology term) | The number of high impact alterations commonly identified in cfDNA and tDNA of each glioblastoma patient |  |  |  |  |  |  |  |  |  |
| --- | --- | --- | --- | --- | --- | --- | --- | --- | --- | --- |
|  | GBM |  | GBM |  | GBM |  | GBM |  | GBM |  |
|  | Patient #GB1 |  | Patient #GB3 |  | Patient #GB5 |  | Patient #GB7 |  | Patient #GB13 |  |
|  | count | % | count | % | count | % | count | % | count | % |
| splice_donor_variant | 1 | 0.0 | 2 | 0.0 | 5 | 0.0 | 2 | 0.0 | 0 | 0.0 |
| splice_acceptor_variant | 0 | 0.0 | 0 | 0.0 | 1 | 0.0 | 1 | 0.0 | 0 | 0.0 |
| stop-gained | 0 | 0.0 | 1 | 0.0 | 1 | 0.0 | 0 | 0.0 | 0 | 0.0 |
| missense_variant | 5 | 0.0 | 2 | 0.0 | 21 | 0.0 | 5 | 0.0 | 1 | 0.0 |
| splice_region_variant | 8 | 0.0 | 9 | 0.0 | 37 | 0.1 | 10 | 0.0 | 6 | 0.0 |
| synonymous_variant | 2 | 0.0 | 3 | 0.0 | 21 | 0.0 | 4 | 0.0 | 5 | 0.0 |
| 5_prime_UTR_variant | 30 | 0.1 | 18 | 0.1 | 148 | 0.2 | 45 | 0.1 | 25 | 0.1 |
| 3_prime_UTR_variant | 244 | 0.7 | 219 | 0.8 | 648 | 0.9 | 293 | 0.9 | 190 | 0.8 |
| non_coding_transcript_exon_variant | 400 | 1.1 | 352 | 1.3 | 1082 | 1.5 | 473 | 1.4 | 290 | 1.3 |
| intron_variant | 16989 | 47.3 | 12707 | 47.5 | 33637 | 47.6 | 15795 | 48.4 | 10780 | 46.5 |
| upstream_gene_variant | 1327 | 3.7 | 1102 | 4.1 | 3417 | 4.8 | 1470 | 4.5 | 997 | 4.3 |
| downstream_gene_variant | 1,254 | 3.5 | 992 | 3.7 | 2778 | 3.9 | 1300 | 4.0 | 824 | 3.6 |
| TF_binding_site_variant | 68 | 0.2 | 62 | 0.2 | 224 | 0.3 | 86 | 0.3 | 65 | 0.3 |
| regulatory_region_variant | 1545 | 4.3 | 1257 | 4.7 | 3702 | 5.2 | 1554 | 4.8 | 1059 | 4.6 |
| intergenic_variant | 14017 | 39.1 | 10000 | 37.4 | 24929 | 35.3 | 11614 | 35.6 | 8954 | 38.6 |

Next, we extended the top-58 selected genes that are commonly mutated in gliomas, by those that were listed in three current studies in cfDNA in glioma^112–114^. Remarkably, for all the patients, the distributions of a number of mutations across the top-58 mutated genes in gliomas were highly conserved (FDR<1%, Fig. 5). This indicates that mutation distribution in cfDNA coincided with glioma type and grade. As a result, of these 58 genes, a total of 34(58%), 38(65%), 20(34%), 27(46%) and 39(67%) genes were identified as mutated in both cfDNA and tDNA of our glioma patients i.e. #GB1, #GB3, #GB5, #GB7 and #GB13, respectively (Table 2). These mutated genes included *TP53*, which encodes a protein that acts as a tumour suppressor in many cancer types including glioma^94^; and the *IDH1*mutation, which correlates negatively with glioma grades II to IV. The *IDH1*mutation is detected at the rates of 77%, 55% and 6%, for grades II, III and IV glioma, respectively^115–120^. Due to these unique characteristics, the *IDH1* mutation serves as a biomarker in the diagnosis and prognosis of glioma ^115,121,122^. These results indicate that our liquid biopsy methodology captures a broad spectrum of known glioma mutations at similar incidence rates as in the tumour biopsies.

**Table 2.**
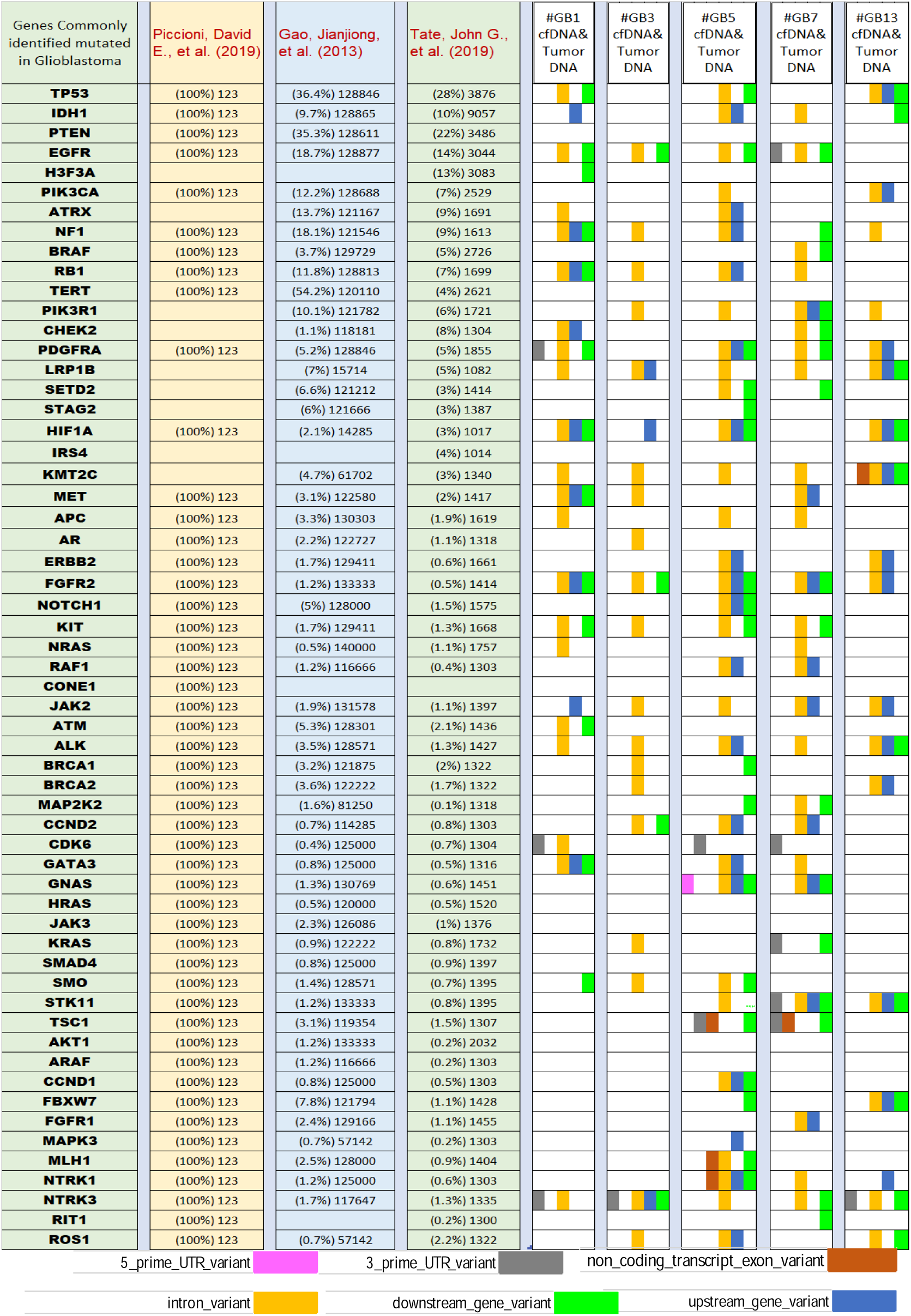
A comparison between high impact mutations identified in five glioblastoma patients (#GB1, #GB3, #GB5, #GB7 and #GB13) and genes reported in the literature as commonly mutated in glioblastoma. Column 1 lists 58 genes that have been found to be commonly mutated in glioblastoma. Columns 2,3 and 4 present data about glioblastoma from 3 major studies (Piccioni, David E., et al. 2019, Gao, Jianjiong, et al. 2013, Tate, John G., et al. 2019). The percentages inside the parentheses indicate the proportions of glioblastoma patients that showed a mutation in corresponding genes in column-1. The numbers outside the parenthesis indicate the total number of glioblastoma patients who were tested in the study. Columns 5,6,7,8 and 9 show the high impact mutations that were identified, commonly in cfDNA and tumour DNA, and in corresponding genes in column 1 in glioblastoma patients #GB1, #GB3, #GB5, #GB7 and #GB13, respectively.

**Fig. 5.**
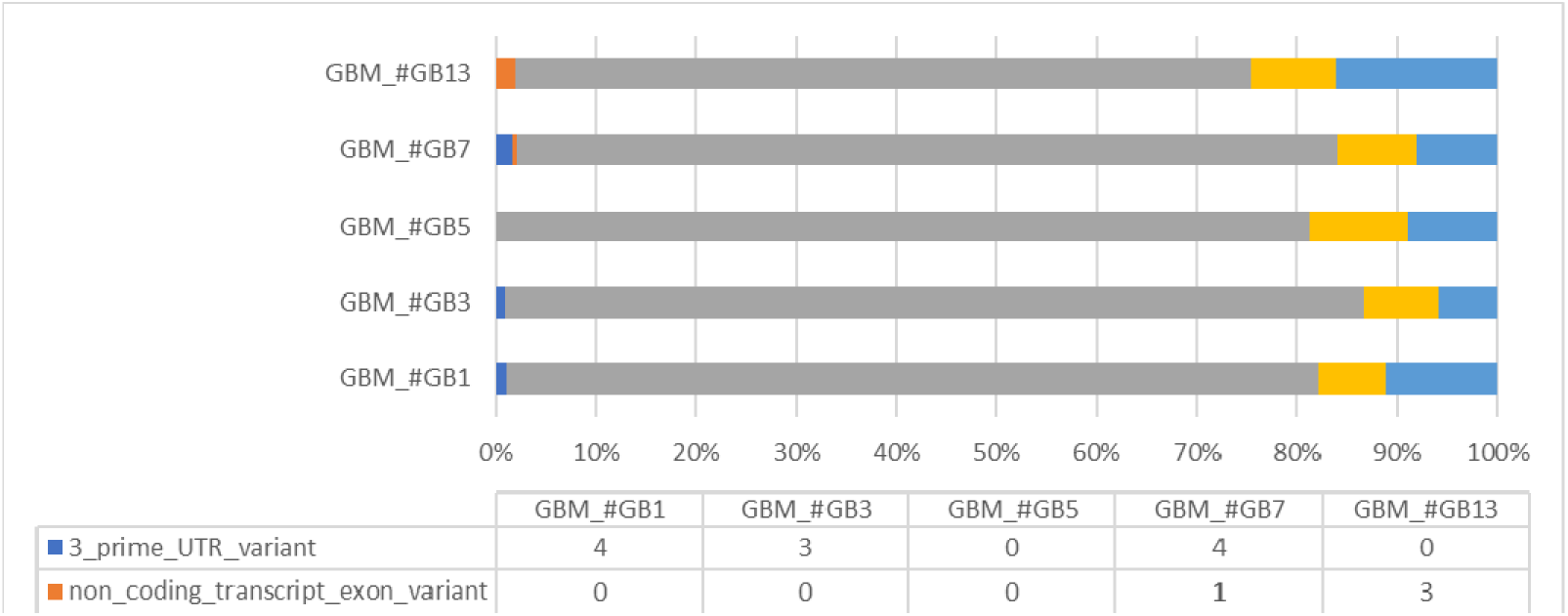
The proportion of high impact alterations in 58 glioma related genes (from Table 2) by 5 consequence types studied in glioblastoma patients #GB1, #GB3, #GB5, #GB7 and #GB13.

### Classification of high-grade gliomas according to mutations in cfDNA

We report a lower level of plasma cfDNA concentrations among low-grade and healthy controls than among high-grade glioma patients. This is presumably due to low numbers of necrotic and apoptotic cells in the former^41,42,44^. This finding suggests a potential sensitivity level for discovering high-grade gliomas using a simple cfDNA concentration test. We examined the possibility of classifying high-grade gliomas using mutations in cfDNA, for the diagnosis of GBM. Accordingly, we selected the 50 most frequently mutated genes from the current study of the mutation landscape in The Cancer Genome Atlas (TCGA) GBM dataset (PanCancer datasets, 291 GBM samples)^123^. Of these 50 genes, 21(42%), 26(52%), 13(26%), 22(44%) and 25(50%) were also mutated in our GBM patients: #GB1, #GB3, #GB5, #GB7 and #GB13, respectively (Table 3). As shown in Table 3, in addition to the most common glioblastoma-related genes, like *IDH1* and *TP53*, we found mutations in the *BRAF* and *EGFR* genes, which are recognized for their involvement in glioma tumour progression^124,125^. These results indicate that mutations found in cfDNA correspond to mutations in tumours, with 95% specificity, such that we were able to distinguish high-grade glioma types of tumours.

**Table 3.**
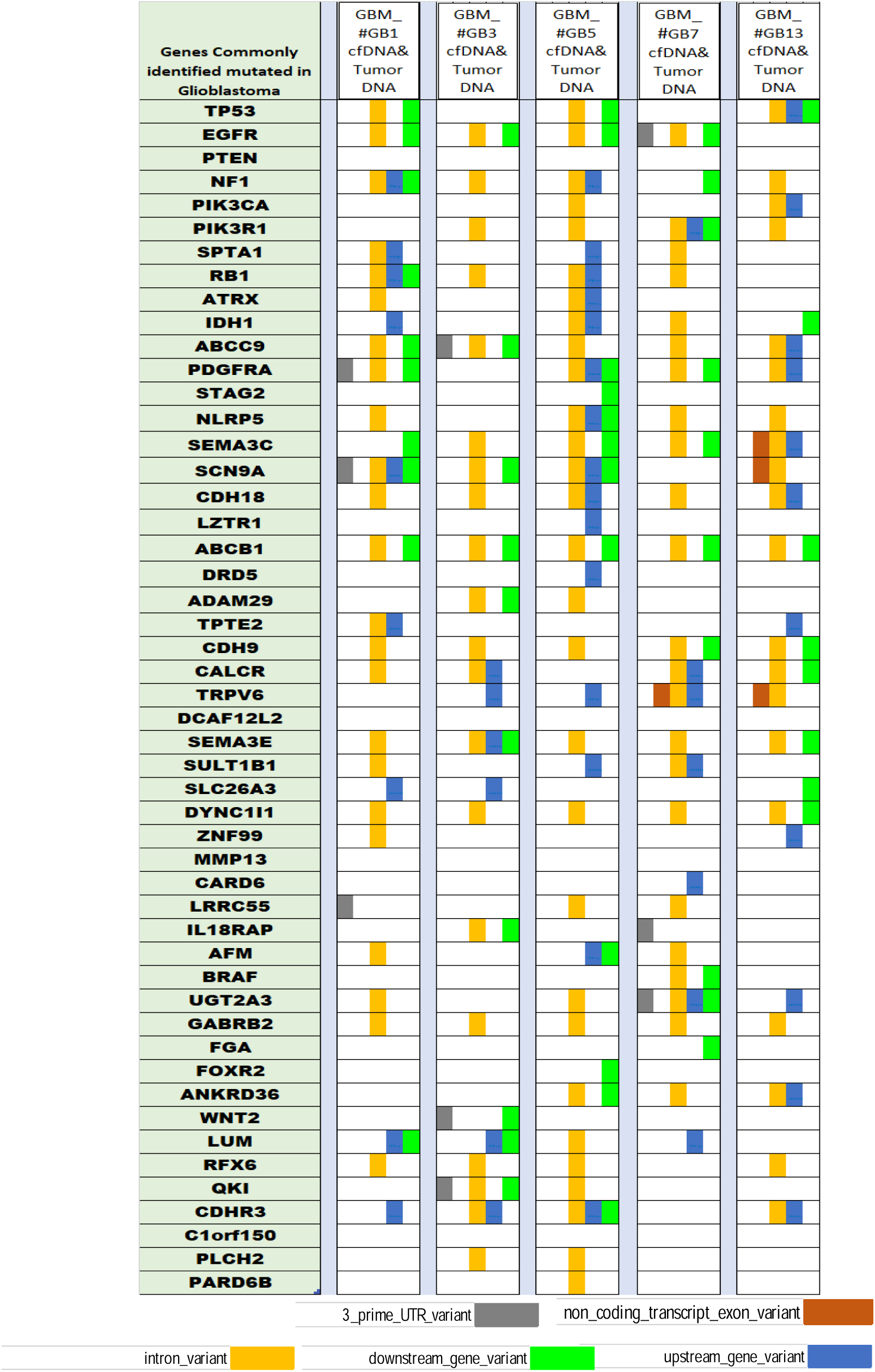
The 50 most frequently mutated genes in 291 glioblastoma patients in the TCGA database, studied by Brennan, Cameron W., et al. (2013) and identified in our five glioblastoma patients (#GB1, #GB3, #GB5, #GB7 and #GB13). Column 1 lists 50 genes that were identified as commonly mutated in 291 glioblastoma patients by Brennan, Cameron W., et al. (2013). Columns 2, 3, 4, 5 and 6 show the high impact mutations that were identified commonly in cfDNA and tumour DNA, and in the corresponding genes in column-1 in glioblastoma patients #GB1, #GB3, #GB5, #GB7 and #GB13, respectively.

Next, we compared the somatic high impact mutations that were commonly shared between cfDNA and tDNA in our five patients to mutation landscape data of GBM from four studies ^112–114,123^ (Tables 2 and 3). We validated these mutations using Sanger sequencing (Fig. 11). Additionally, we found that cfDNA produces a high-level profiling of somatic mutations in all GBM patients. Particularly, we found mutations in genes that are strongly involved in GBM, i.e. *EGFR* (3’ UTR variant, intron variant, downstream gene variant), *IDH1* (upstream gene variant, intron variant, downstream gene variant), *PDGFRA* (3’ UTR variant, intron variant, downstream gene variant, upstream gene variant), *PIK3CA* (intron variant, upstream gene variant), *PIK3R1* (upstream gene variant, downstream gene variant) and *TP53* (upstream gene variant, intron variant, downstream gene variant). Finally, we found that tumour-suppressors were mostly removed from the gliomas by **missense mutations** (Table S3, Supplementary Material) and that oncogenes were gained in the annotated data of our mapped **cfDNA sequences** (Table S3). These results indicate that liquid biopsy may uncover particular somatic mutations that differentiate glioma patients from patients with other tumour types.

**Fig. 6.**
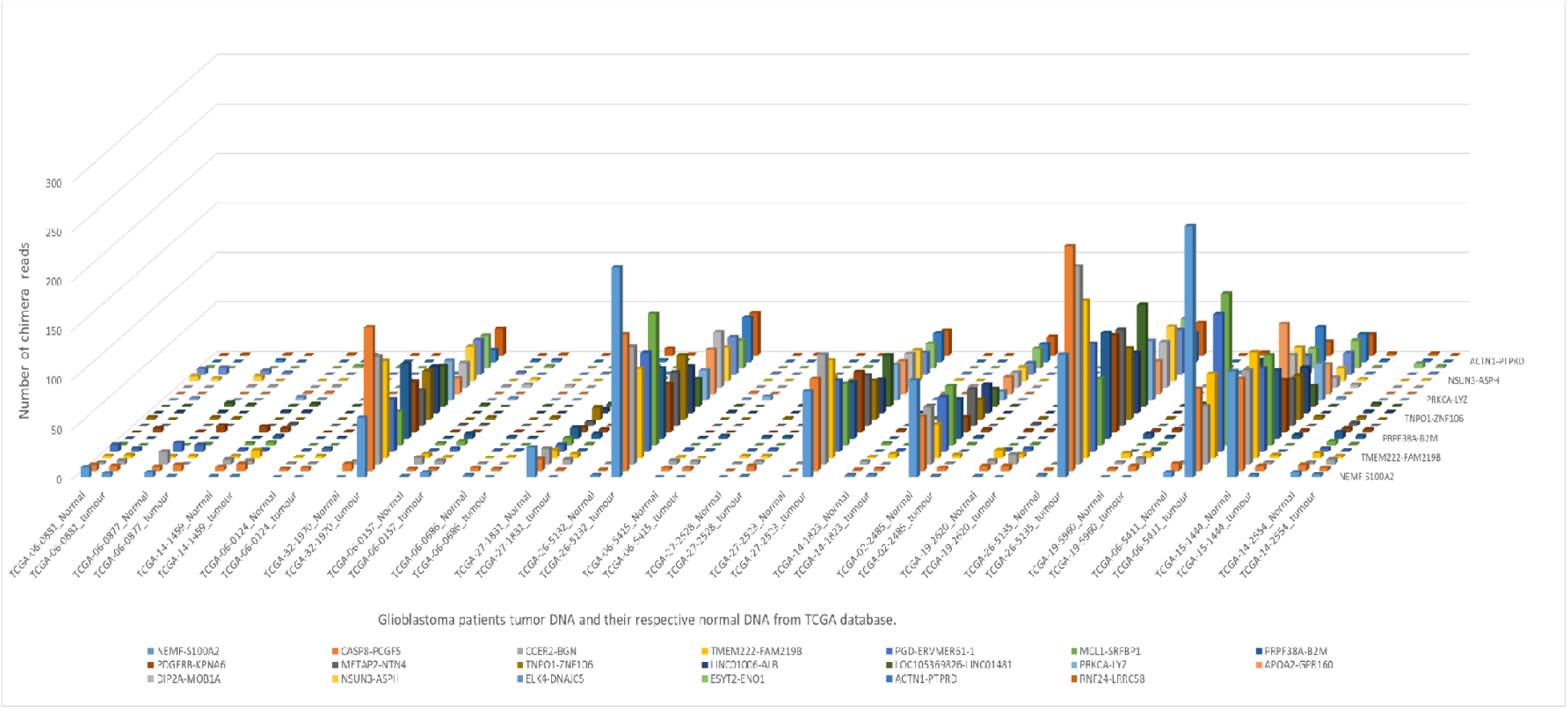
Frequently occurring fusion genes studied in DNA, tumour DNA and respective germline DNA of 20 glioblastoma patients, archived from The Cancer Genome Atlas database.

**Fig. 7.**
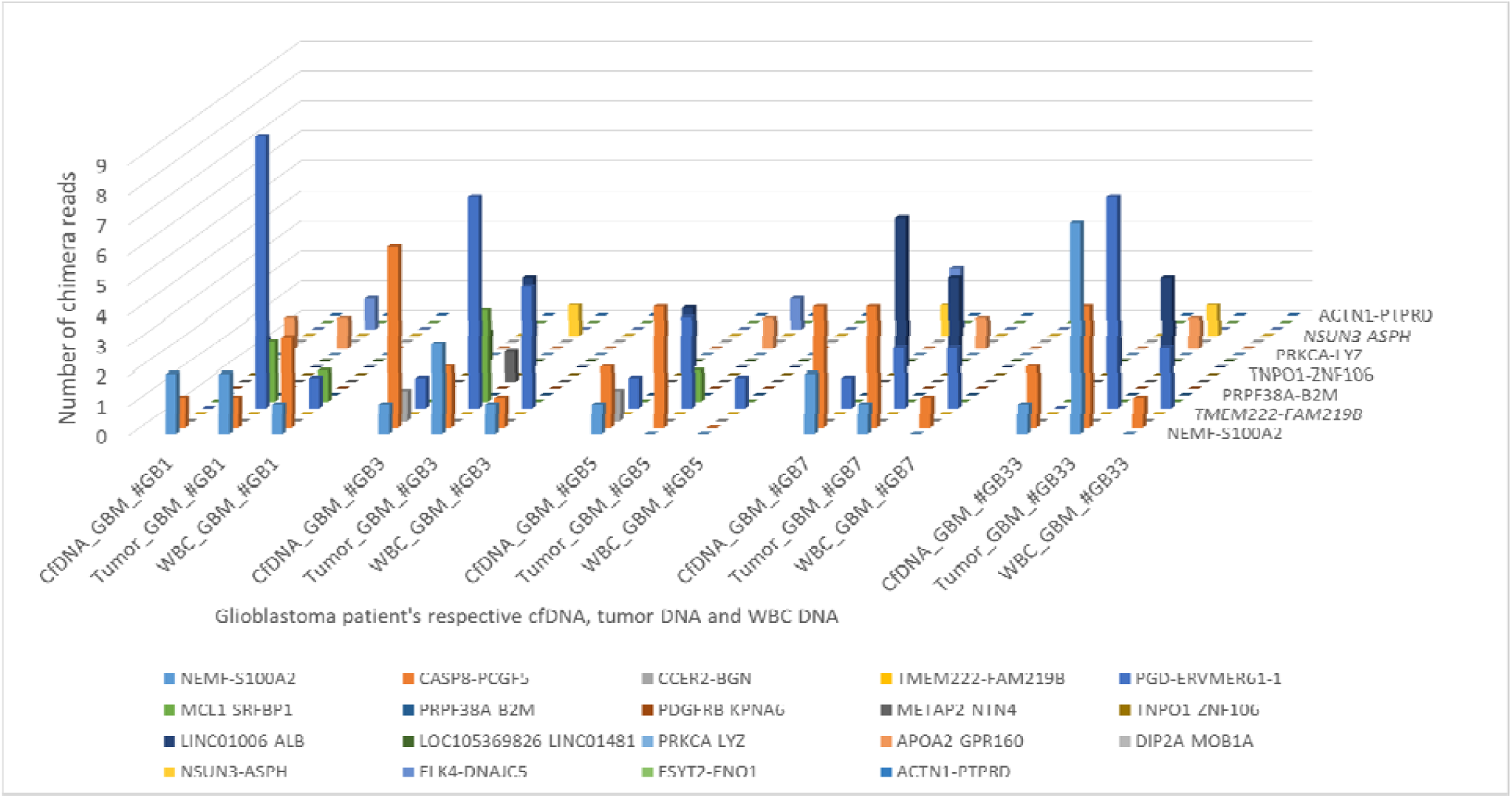
Twenty fusion genes studied in glioblastoma patients from The Cancer Genome Atlas dabtase (figure 5); and also analysed in cell-free DNA, tumour DNA and germline DNA of glioblastoma patients #GB1, #GB3, #GB5, #GB7 and #GB13.

**Fig. 8.**
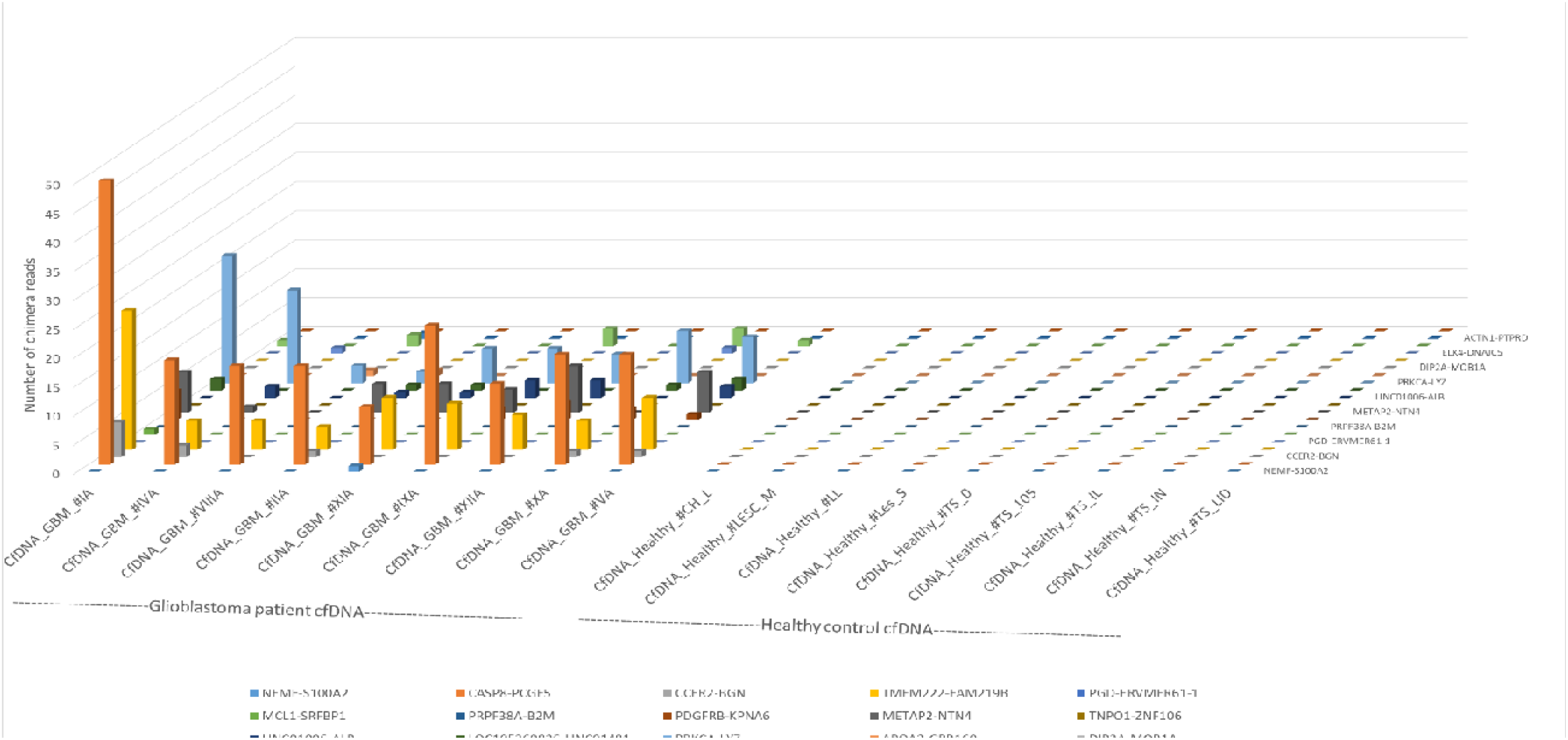
Twenty fusion genes studied in glioblastoma patients from The Cancer Genome Atlas database (figure 5), and also analysed in cell-free DNA of 9 glioblastoma patients and 9 healthy controls.

**Fig. 9.**
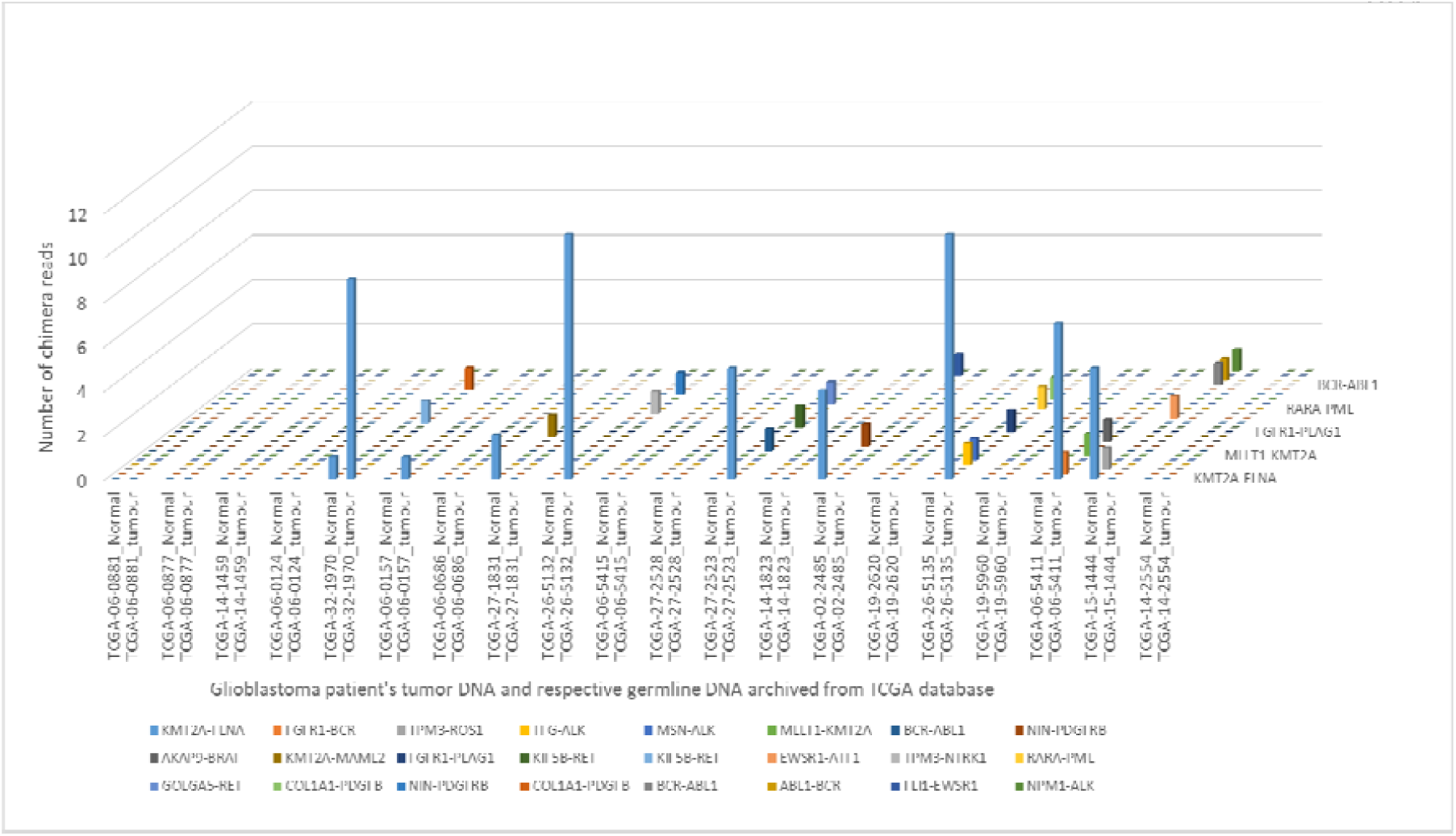
Druggable fusion genes identified in glioblastoma samples archived from The Cancer Genome Atlas database.

**Fig. 10.**
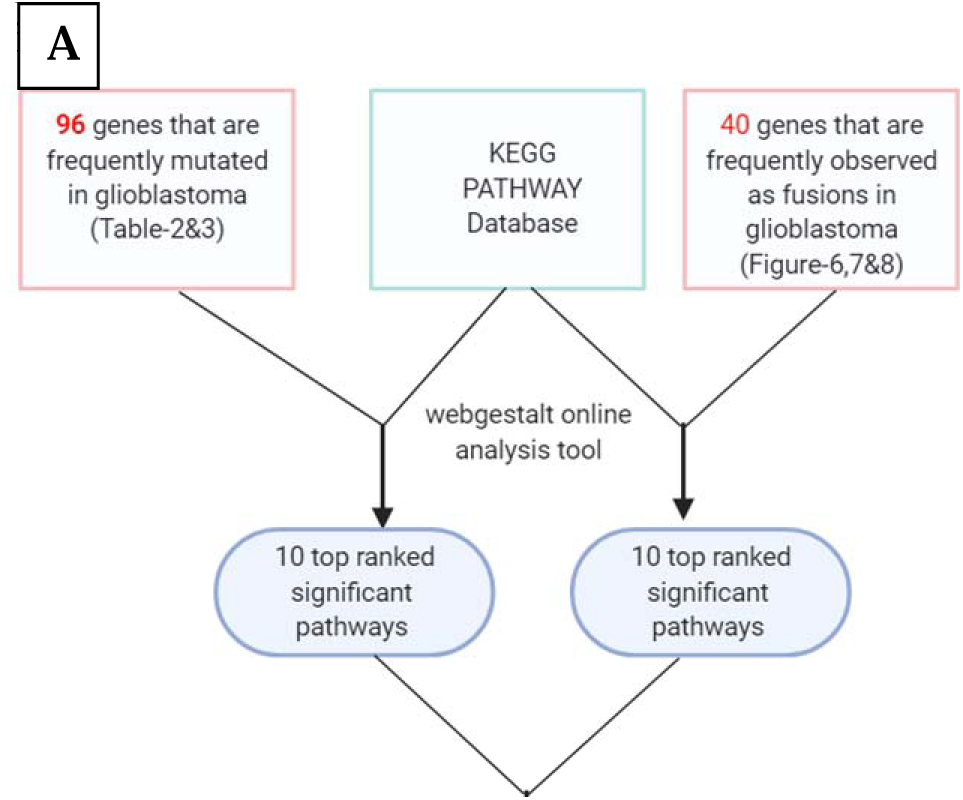
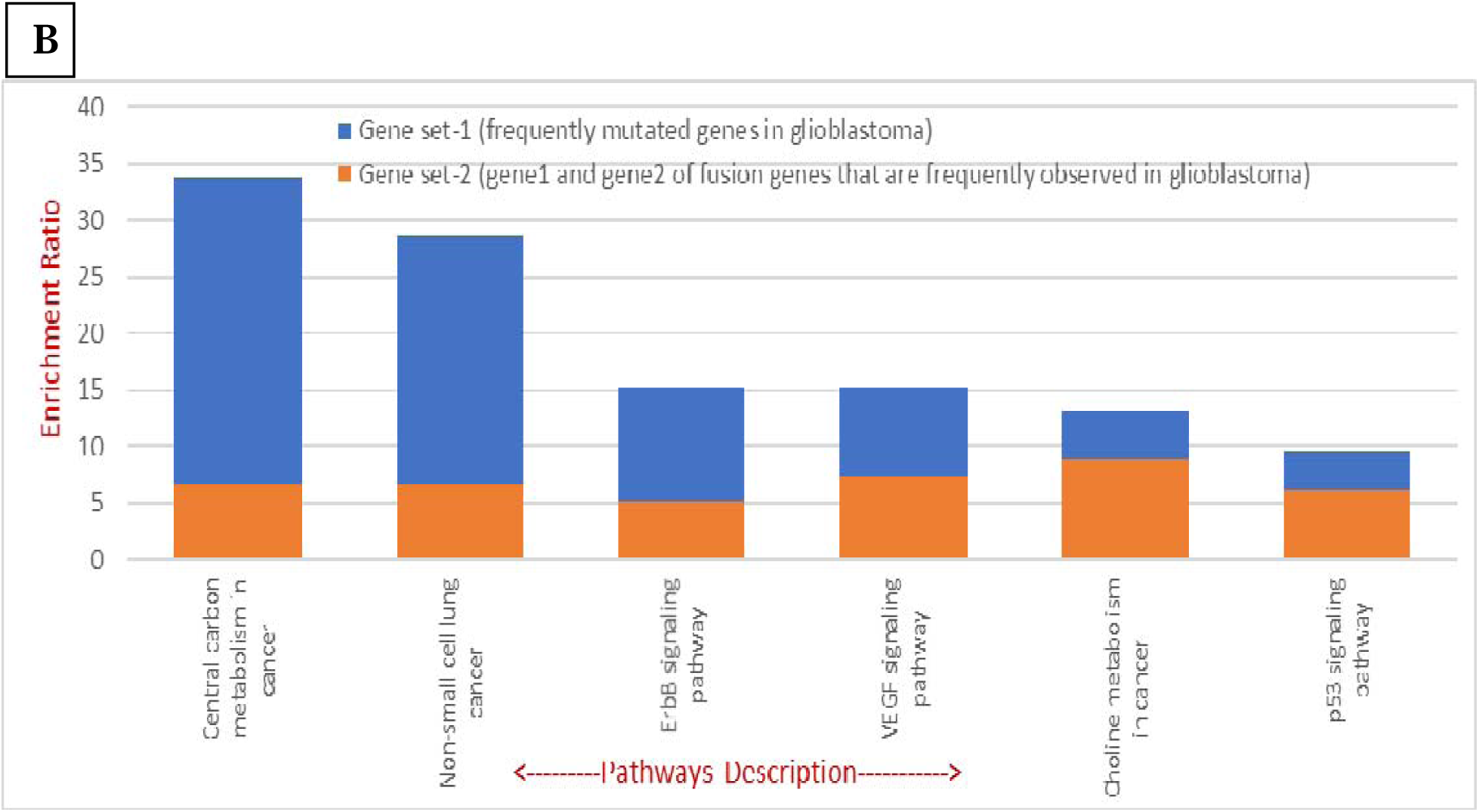
Gene set enrichment analysis study. **A**. Gene set enrichment analysis flowchart. A total of 96 genes that were identified as frequently mutated in glioblastoma (from tables 2&3); and 40 genes that are frequently observed as fusions in glioblastoma (from figures 6,7&8) were analysed against human pathway database KEGG, using online analysis tool webgestalt **B**. The bar graph shows 6 significant pathways, for which at least 1 gene was involved from both: frequently mutated glioblastoma genes and genes that were identified as frequent fusions in glioblastoma.

**Fig. 11.**
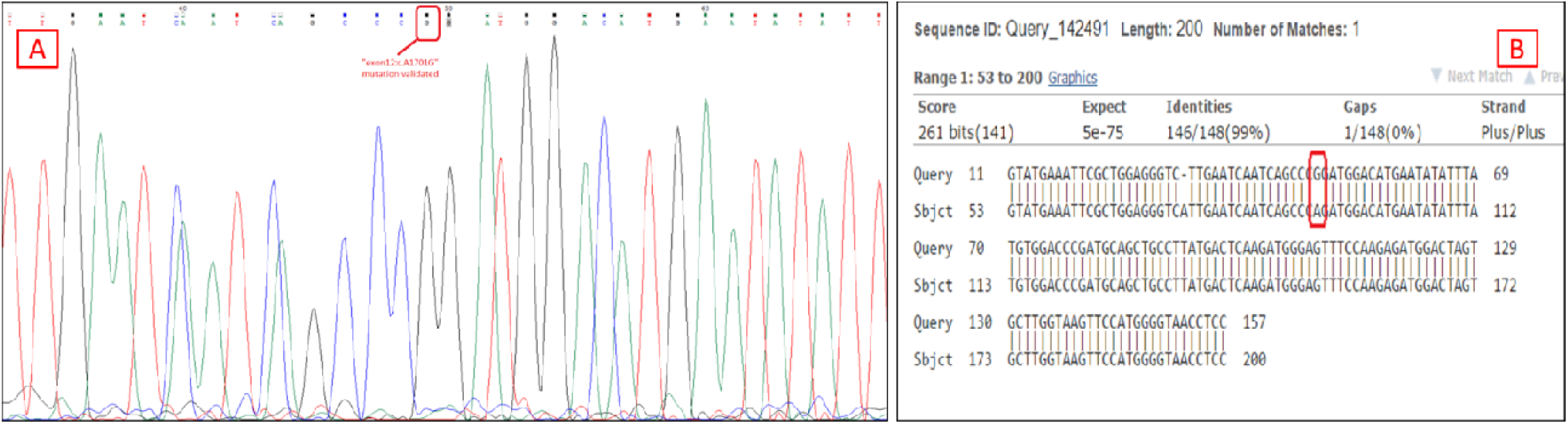
Mutation validation by Sanger sequencing. PDGFRA mutation ‘exon12:c.A1701G’ validation using Sanger sequencing, **A-**Sanger sequence visualization in Chromas^®^ software, **B-**BLAST result of the Sanger sequence when compared with reference human genome hg38.

### Fusion gene analysis

We hypothesized that fusion genes may contribute to glioma tumour formation, in addition to the point mutations described above, and that specific fusion vs. mutation combinations may be unique to glioma. We analysed DNA sequences from 20 normal samples, 18 of 20 glioma samples and seven tDNA and WBC gDNA of TCGA GBM patients. We searched for fusions, using our ChiTaRS 5.0 reference database (http://chitars.md.biu.ac.il/)^126^. Fig. 6 shows the top twenty gene fusions identified in tDNA, but not in the gDNA of TCGA GBM patients #321970, #265132, #272523, #265135 and #065411. On the contrary, in TCGA GBM patients #022485 and #151444, these top twenty gene fusions were identified in gDNA but not in their respective tDNA. Comparing these twenty fusions with cfDNA, tDNA and gDNA from five GBM patients identified four unique gene fusions, i.e. APOA2-GPR160 (#GB1), CASP8-PCGF5 (#GB5), NEMF-S100A2 (#GB7, #GB13) and LINC01006-ALB (#GB7, #GB13) in cfDNA and tDNA, but not in the respective gDNA (Fig. 7). Next, we compared cfDNA of nine GBM patients and nine healthy controls (Fig. 8). Interestingly, the fusions that were identified in the cfDNA of the nine GBM patients were not identified in the cfDNA of any of the nine healthy controls. These results indicate that a fusion gene signature may be readily detectable in glioma patients, thus distinguishing them from non-cancer controls, with high specificity and sensitivity (at 1% FDR). Furthermore, we analysed our datasets to identify hits among predicted 1207 druggable fusions that had been collected in the ChiTaRS 5.0 database and characterized by a preserved tyrosine kinase domain targeted by chemotherapy drugs. We identified 24 druggable fusions for the crizotinib analogues (i.e. entrectinib and larotrectinib) in tumours of TCGA GBM patients and in gDNA (Fig. 9, Table 4). Particularly, we found two druggable fusion genes in the gDNA of GBM patients #GB3 and #GB7 (Table 4), two druggable fusion genes in the cfDNA of GBM patients #IA and #VIIIA (Table 4), and one druggable fusion gene in the cfDNA of the healthy control #TS_0 (Table 4). These results indicate that some glioma patients have druggable biomarkers for the entrectinib or larotrectinib drug (Table 4), which may be used for personalized and improved chemo-radiotherapy protocols. However, crizotinib has poor penetration into the CNS and, therefore, is not a good drug candidate for treating brain tumours^127,128^. Therefore, we considered new drugs to treat NTRK and also ROS1 fusions that have shown some brain tumour activities (data are scarce, mainly from brain metastases) like entrectinib and larotrectinib^129^. Regarding ALK inhibitors there are also new drugs with better brain penetration such as lorlatinib^130^ and brigatinib^131^.

**Table 4.**
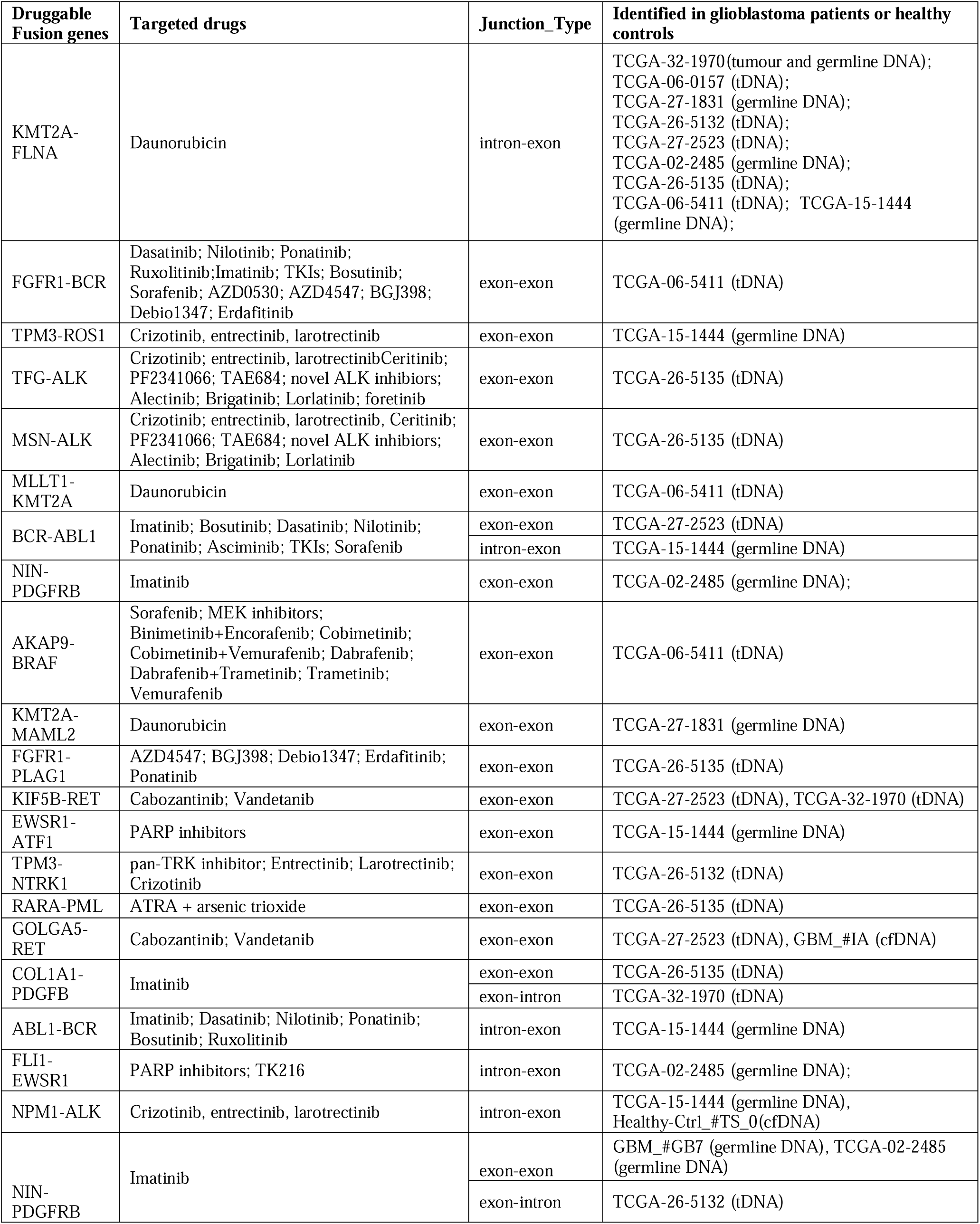
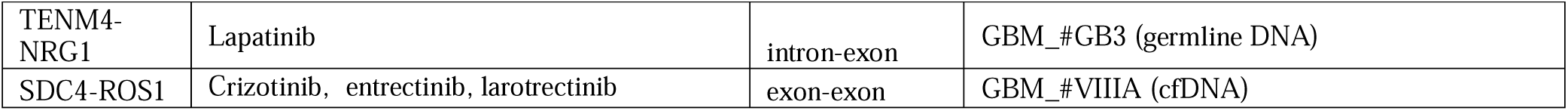
Druggable fusion genes and their targeted drugs identified in glioblastoma samples archived from The Cancer Genome Atlas (TCGA) database; cell-free DNA (cfDNA), tumour DNA (tDNA) and germline DNA of glioblastoma patients, and cfDNA of healthy controls.

### Gene enrichment analysis

Since functional mutations and fusions act to disrupt key metabolic pathways in cancer cells, we examined whether glioma-specific pathway disruptions could potentially be treated with targeted drug combinations. First, we found that a specific subset of fusions (n=5) incorporates a druggable oncogene target that is likely to respond to crizotinib analogues i.e. entrectinib and larotrectinib. Furthermore, 15 additional fusions found in 4 glioma patients and 9 glioma samples archived from the TCGA database indicated potential druggability of 38.2% of all patient samples analysed, via 40 identified genes that are frequently involved in gene fusions (Fig. 5,6 &7). Second, we hypothesized that particular pathways in gliomas were affected by mutations, as well as by fusions. We analysed the gene set and identified pathway enrichment for 96 genes that were previously reported as frequently mutated in glioma patients^112,114,123^. Additionally, we analysed another gene set, which included 40 genes that were frequently observed in **fusions** in glioma patients. The KEGG PATHWAY database was used for the analysis, and the 10 most significant pathways were identified from each gene set (Fig.10A). The significant pathways for each gene set were then compared between each other. Six significant pathways, namely, the ErbB signaling pathway, the VEGF signaling pathway, the choline metabolism pathway, central carbon metabolism in cancer, the p53 signalling pathway and pathways in non-small cell lung cancer were identified as common between these two gene sets. Such analysis shows that cancer specific pathways are similar and targeted by either **acquiring gene mutations or by forming gene fusions**. Thus, a comprehensive study of both gene mutations and fusions can contribute to the understanding of targeted pathways in glioma patients.

## Discussion

In this study, we showed that cfDNA concentration in the plasma of GBM patients is higher than in low-grade glioma patients, and higher than in healthy persons. The direct association of cfDNA concentration and tDNA burden in plasma was previously reported in a few studies^32,45–47^. Moreover, cfDNA concentration was shown to be a prognostic biomarker in colorectal, ovarian and breast cancers, non-small cell lung cancer (NSCLC) and melanoma cancer^64,132–134^. Therefore, cfDNA concentration can serve as a potential combined biomarker in glioma liquid biopsy, for the diagnosis, prognosis and prediction of glioma tumours. We found that tDNA fragments are continuously circulating in the plasma of GBM patients; this probably reflects the release of more tDNA into the circulation. These findings concur with previous reports^135,136^. The implication is that tumour cfDNA can be used for liquid biopsy tests in GBM patients. We extended these findings by the novel fusions and, particularly, the druggable fusion targets for crizotinib and its new analogues, namely entrectinib and larotrectinib. This demonstrates enhanced CNS penetration in glioma patients, as an alternative line of personalized treatment.

Several studies have reported the dynamics of cfDNA based mutations in patients with various cancers. Early detection of these mutations can be helpful in cancer diagnosis, and in determining treatment response outcome ahead of standard methods. This will contribute to predicting disease state and cancer prognosis^64,132– 134^. However, challenges still remain, as distinct diagnostic biomarkers with high sensitivity and specificity have not been identified for most cancer types, including glioma subtypes. In this study, we added to mutation analysis, gene-gene fusions that are known for their tissue specificity, as well as for cancer specificity^86,91^. We describe outcomes of mutation analysis of cfDNA and tDNA in glioma patients, based on previous glioma mutation landscapes. This analysis classifies glioma types and provides information on the genes that were mutated and the pathways affected by the mutations. Mutations in genes such as *IDH1/2, TERT, BRAF, EGFR* and *ATRX* have been studied for their associations with prognosis and with treatment response^11,115–120,137,138,139,140^.

Particularly, *IDH* is a major prognostic feature for astrocytomas. Even in tumours that do not show classical histologic features of GBM, *IDH* wild type status is closely linked to lower overall survival and rapid deterioration^141^. Another important mutation found in our glioma samples is the *TERT* promoter mutation, which is observed in 70% of oligodendroglioma and in about 70–80% of primary GBM. This mutation occurs mainly in the 124bp-146bp upstream of the transcription start site^142^. The *TERT* promoter mutation has been shown to be a typical feature in progressive GBM^137,138^. Further, the *TERT* promoter mutation was found to be associated with shorter overall survival and to be negatively correlated with the grade of astrocytoma^11^. A positive correlation with *EGFR* amplification and negative correlations with *IDH1* mutations are key features of *TERT* promoter mutations in GBM tumours^11^. This suggests that these mutations may be important prognostic markers for patients with glioma. Moreover, we identified a mutation in the *ATRX* gene that occurs exclusively in astrocytomas^139,140^. Thus, the *ATRX* mutation is useful in glioma tumour diagnosis, since the presence of both *IDH* and *ATRX* mutations is considered a reliable indication of lower grade astrocytomas^139,140^. In contrast, the *TERT* promoter mutation is seen mainly in primary GBM and oligodendroglioma^78,94^. The *BRAF* gene encodes the BRAF protein. This serine/threonine kinase serves as an immediate downstream effector of the MAPK signalling cascade, a signal transduction pathway that modulates cell proliferation and survival. *BRAF* alterations leading to MAPK pathway activation have been identified in gliomas and glio-neuronal tumours of the CNS^124^. The *EGFR* mutation is another important gene mutation that we identified. Thus, enhanced activation of *EGFR* can occur through a variety of mechanisms, both ligand-dependent and ligand-independent^125^. In particular, substantial evidence suggests that *EGFR* is overexpressed in most primary GBM and in some secondary GBMs, and is characteristic of more aggressive GBM phenotypes^125^. Moreover, we found mutations in a set of genes (i.e., *EGFR, IDH1, PDGFRA, PIK3CA, PIK3R1* and *TP53*) that are involved in the central carbon metabolism pathway in cancer^143^. This pathway helps cancer cells consume a large amount of glucose to maintain a high rate of glycolysis, and provides intermediate molecules to synthesize most of the macromolecules required for the duplication of cancer cell biomass and the genome^144^. Therefore, cfDNA testing may serve as an effective liquid biopsy platform in low- and high-grade gliomas and, particularly, in GBM patients.

We detected the abovementioned mutations in the cfDNA and tDNA of GBM patients. This suggests that liquid biopsy can provide molecular signatures for glioma management. Moreover, these molecular signatures can be monitored serially since liquid biopsy requires only a simple blood test and can be done frequently during the disease course. In 20 TCGA archived GBM patients and 14 of our GBM patients, we found that 20 fusion genes **were either formed in large numbers of cfDNA molecules or were rarely formed**. The phenomenon of this activity of fusion genes is still not understood. However, the complete absence of these fusion genes in a healthy control cohort supports the disease specificity of the fusion genes. Next, we showed by gene enrichment analysis that these frequently observed fusion genes and the 96 most frequent genes from the glioma mutation landscape share common pathways that are significant in gliomas. These include the ErbB signaling pathway, the VEGF signaling pathway, the choline metabolism pathway, the central carbon metabolism pathway and the p53 signalling pathway. The ErbB signaling pathway is enriched for both mutations and fusions in gliomas. Receptor proteins ErbB1, ErbB2, ErbB3 and ErbB4 belong to the ErbB receptor family of tyrosine kinases. Upon ligand induction, the receptor activates downstream signaling pathways that lead to cell migration, cell proliferation and anti-apoptosis processes. Mutations in these receptors lead to constitutive activation of receptors, independent of ligand induction. This results in increased cell migration, cell proliferation and anti-apoptosis processes. This alteration is recognized as a key target for alteration in many other tumour types^145^. The VEGF signaling pathway (enriched for both mutations and fusions in gliomas) activates angiogenic protein VEGF under hypoxic conditions and increases vascular permeability. GBM tumours are often hypoxic and require increased angiogenesis. This hypoxia triggers VEGF overexpression, and this contributes to the irregular vasculature associated with GBM^146^. The choline metabolism pathway was also targeted by mutations and fusions. This pathway is characterized by increased phosphocholine and total choline-containing compounds. Abnormal choline metabolism is influenced by hypoxia conditions in the tumour microenvironment and is found to cause the transformation of non-malignant cells to malignant cells^147^. Thus, gene set enrichment analysis can compare two sets: genes frequently identified as mutated in GBM and genes that were frequently identified as fusions in GBM in the current study. This indicates that fusion genes, together with mutations, directly target the disease-causing pathways in glioma tumours. Therefore, fusion genes can be studied together with mutations for their combined use in estimating precise prognosis and monitoring treatment response by cfDNA in primary brain tumours, and specifically in gliomas.

In addition to the role of liquid biopsy in estimating prognosis and treatment response, this technique carries the potential to improve glioma management by providing druggable fusion targets for treating patients. Using liquid biopsy, we identified 15 druggable fusions in four glioma patients. Nine TCGA archived glioma samples indicated potential druggability of 38.2% of all the patient samples analysed. In three GBM patients, we identified *ALK-*based druggable fusions, namely *TFG-ALK, MSN-ALK* and *NPM1-ALK*. In a previous study, NSCLC patients with *ALK*-positive fusions were treated with the kinase inhibitor crizotinib for a mean duration of 6.4 months. The overall response rate was 57% (47 of 82 patients: 46 confirmed partial responses and 1 confirmed complete response); 27 patients (33%) had stable disease^96,148^. Two additional important druggable fusions that we found in two GBM patients were *SDC4-ROS1* and *TPM3-ROS1. ROS1-*based fusions were initially discovered in the human glioblastoma cell line U118MG^149,150^. A previous *in-vitro* study showed that treatment with ROS1 inhibitor crizotinib was anti-proliferative and that it downregulated signalling pathways that are critical for growth and survival^151^. We detected *BCR-ABL1* fusions in two, *NIN-PDGFRB* in three and *COL1A1-PDGFB* in one of our GBM patients. These three fusion genes can be targeted by the common drug imatinib (Gleevec)^152,153,154^. Therefore, targeted drugs with improved brain penetration should be tested, based on the dynamics of fusions detected in patients’ plasma.

In conclusion, we showed that liquid biopsy may have an important role in glioma management. This is due to its non-invasive nature and its ability to provide broad range information on real-time activity of mutations and gene fusions in brain tumour patients. Specific gene mutations and fusion genes can act as combined markers for estimating prognosis and treatment outcomes in these patients. Therapeutic druggable fusion gene targets can be identified using liquid biopsy; this will contribute to precise treatment of glioma patients using non-invasive liquid biopsy diagnostic technique.

## Materials & methods

### Sample collection, storage and maintenance

From 27 glioma patients, brain tumour samples (fresh frozen), blood plasma and peripheral blood mononuclear cells (PBMCs) were obtained from several hospitals and from a biorepository. Three samples were provided by Prof. Rainer Glaβ, the Department of Neurosurgery, Ludwig-Maximilians-University, Munich, Germany; 9 samples were provided by Dr. Charlotte Flueh, the Department of Neurosurgery, University Hospital of Schleswig-Holstein, Campus Kiel, Kiel, Germany, 10 samples were provided by Prof. Tali Siegal, Neuro-Oncology Center, Rabin Medical Center, Petach Tikva, Israel, and 5 samples were provided by The Israeli Biorepository Network for Research (MIDGAM). From 14 healthy controls, we collected blood samples that were separated into plasma and PBMCs. Blood was collected into EDTA-anticoagulated tubes, and plasma was separated within 2 hours of collection. About 1-2 ml of plasma and about 1 ml of PBMC were separated from each blood sample. Both samples were kept at -80° C and were shipped on dry ice.

### DNA isolation

Cell-free DNA (cfDNA) was isolated using QIAamp Circulating Nucleic Acid Kit (Qiagen^®^, Germany) from different volumes of plasma samples (850µl to 2ml), and from 5ml culture media collected from glioblastoma cell line cultures. All samples were processed according to the manufacturer’s standard protocol. NucleoSpin^®^ Tissue kit (Macherey-Nagel, Germany) was used to process genomic DNA from 25 mg of brain tumour biopsies and from 0.5ml of PBMC samples from each patient. Isolated DNA samples were stored at -20°C until further use.

### Nucleosomal DNA isolation

Nucleosomal DNA was isolated from the glioblastoma cell line LN-229 and astrocytoma grade-IV cell line CCF-STTG1 using ACTIVE MOTIF^®^ Nucleosome Preparation Kit™. Isolated DNA samples were then stored at -20°C until their further use.

### DNA quantification

All isolated DNA samples were quantified by Qubit^®^ dsDNA HS assay using Qubit^®^ 2.0 fluorometer. The assay was performed according to the manufacturer’s standard protocol. Fluorescence was measured at 485/530 nm on a Qubit^®^ 2.0 fluorometer to determine DNA concentration for each sample. A Bioanalyzer 2100 DNA High Sensitivity assay was performed to estimate the fragment size distribution of isolated cfDNA samples.

### Next-generation sequencing (NGS) and data analysis

NEBNext^®^ Ultra™ II DNA Library Prep Kit was used for NGS library preparation and sample libraries were sequenced on Illumina HiSeq 2500 and Illumina NextSeq 550 platforms. The covaris fragmentation step was performed only for tDNA and germline DNA from PBMCs, and not for cfDNA.

NGS libraries were prepared for cfDNA, tDNA and germline DNA (PBMC) samples, and were successfully sequenced on Illumina HiSeq2500 and Illumina NextSeq550 platforms. Samples from GBM patients were sequenced as whole genome sequencing at an average of 5x coverage (cfDNA and tDNA) and whole exome sequencing at an average of 180X coverage (for germline DNA).

NGS data were subject to quality control analysis of raw sequencing reads using FastQC and an additional in-house shell script. Adapters and low-quality sequences were trimmed using Cutadapt tool^155^. Remaining reads were mapped to a human genome reference (hg38) using Bowtie2^156^ and SAMtools^157^. Next, SNVs were identified using bcftools^158^ mpileup for each sample. Further, germline variants identified in PBMC DNA were removed from respective patients’ tumours and cfDNA variants, and considered as somatic variants. Somatic variants from cfDNA and tDNA were annotated using standalone Ensembl Variant Effect Predictor (VEP) pipeline^159^.

Reads unmapped to the reference human genome (hg38) were extracted using SAMtools^157^. These were mapped against the reference database of unique chimera junction sequences, ChiTaRS-3.1^110^, using an in-house chimera algorithm.

### Gene set enrichment analysis

Gene set enrichment analysis was performed using the online tool ‘webgestalt’^160^, in which two gene sets (i.e., a set of genes commonly mutated in GBM and a set of genes that fused with high frequencies in GBM tumours and cfDNA) are analyzed against the KEGG PATHWAY^161,162^ database. The aim is identification of the 100 most significant pathways connected to the genes in each gene set. Significant pathways of each gene set were further compared to identify common pathways between the sets (Fig.-11A).

### Mutation validation using Sanger sequencing

Twenty-two point mutations from tumours and cfDNA of GBM patients were selected for validation of Sanger sequencing. Primers were designed using Primer3 (v. 0.4.0)^163^. All amplified PCR products were isolated using silica membrane spin column technique (NucleoSpin^®^ Gel, PCR clean up kit Macherey-Nagel, Germany) and were eluted in 20 µl of nuclease free water. PCR products were then processed for Sanger sequencing and the results were analysed using Basic Local Alignment Search Tool (BLAST^®^)^164^ and Chromas^®^ 2.6.2^165^.

