## Supplementary-2 for "A liquid biopsy platform for detecting gene-gene fusions as glioma diagnostic biomarkers and drug targets"

**1. The Azrieli Faculty of Medicine, Bar-Ilan University, 1311502, Safed, Israel.**

**2. Department of Neurosurgery, Ludwig-Maximilians-University, 81377, Munich, Germany.**

**3. Department of Neurosurgery, University Hospital of Schleswig-Holstein, Campus Kiel, 24105, Kiel, Germany.**

**4. Neuro-Oncology Center, Rabin Medical Center, 4941492, Petach Tikva, Israel.**

*To whom correspondence should be addressed.

| Sample ID | Sample type | Disease/Healthy control | Bioanalyzer/Tapestation |
|---|---|---|---|
| #843 | Cell-free DNA | Glioblastoma | 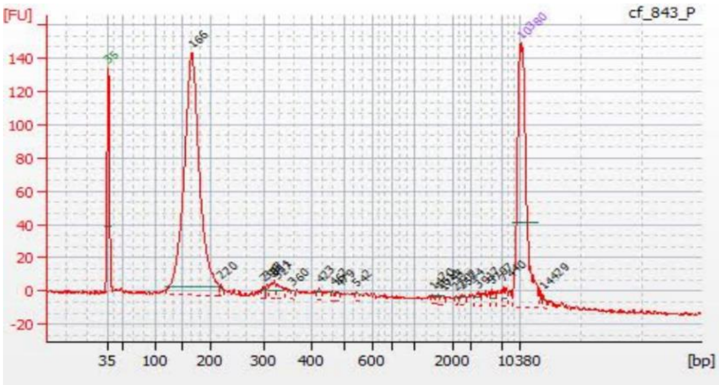 |
| #GB1 | Cell-free DNA | Glioblastoma | 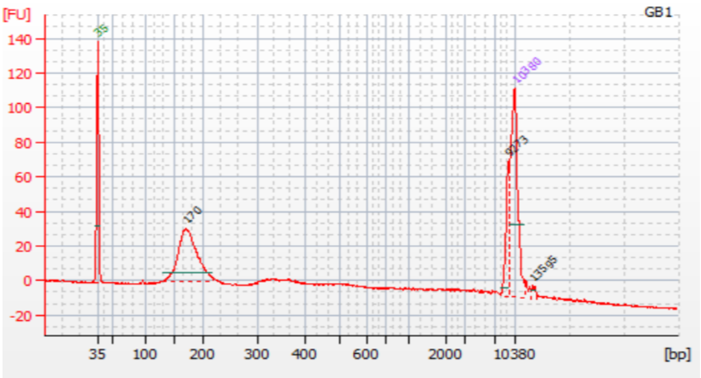 |

<table>
<tr><td>#GB3</td><td>Cell-free DNA</td><td>Glioblastoma</td><td>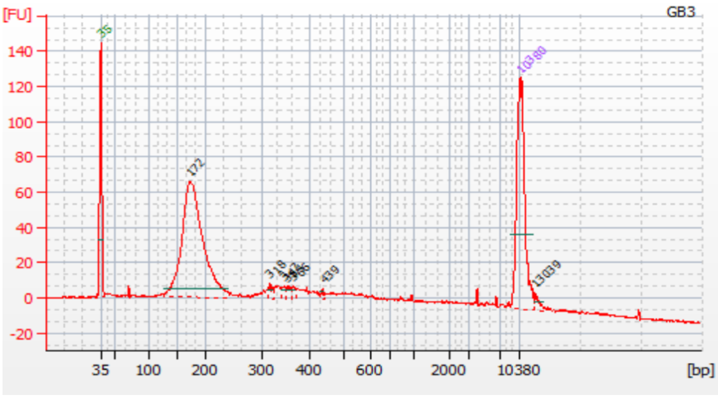
</td></tr>
<tr><td>#GB5</td><td>Cell-free DNA</td><td>Glioblastoma</td><td>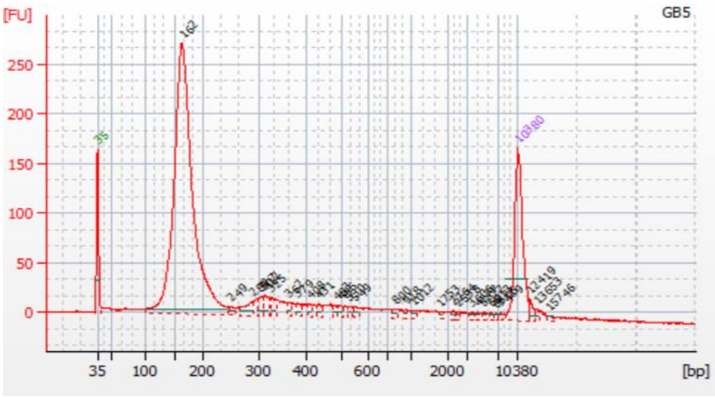
</td></tr>
<tr><td>#GB7</td><td>Cell-free DNA</td><td>Glioblastoma</td><td>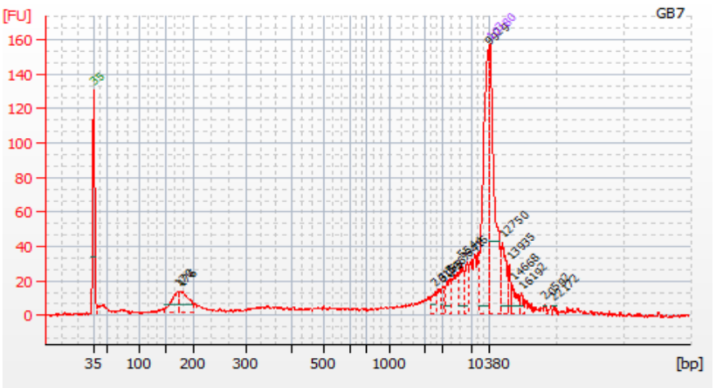
</td></tr>
</table>

| | | | |
|---|---|---|---|
| #GB13 | Cell-free DNA | Glioblastoma | 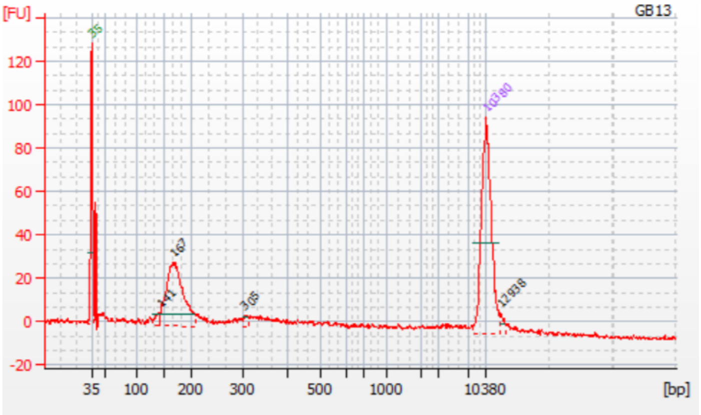 |
| #I | Cell-free DNA | Glioblastoma | 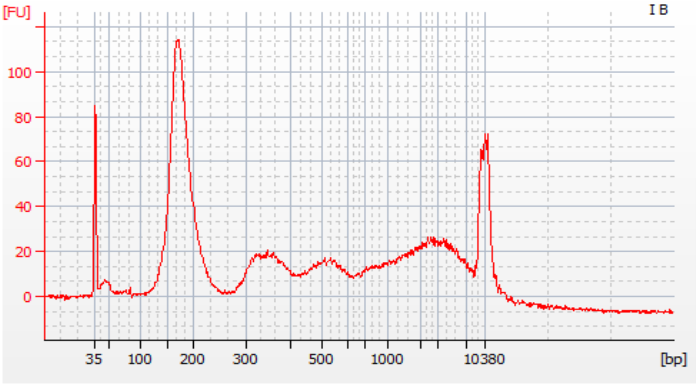 |
| #II | Cell-free DNA | Glioblastoma | 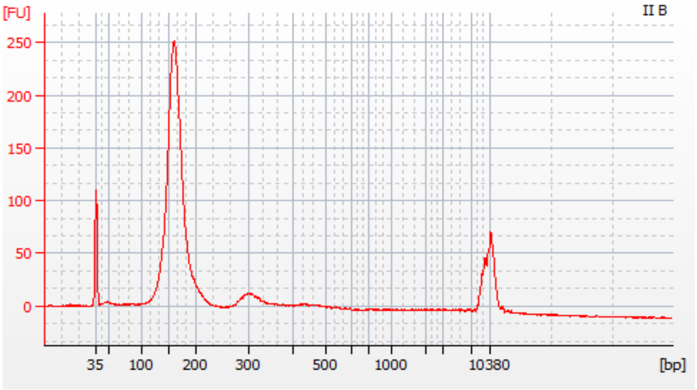 |

| | | | |
|---|---|---|---|
| #IV | Cell-free DNA | Glioblastoma | 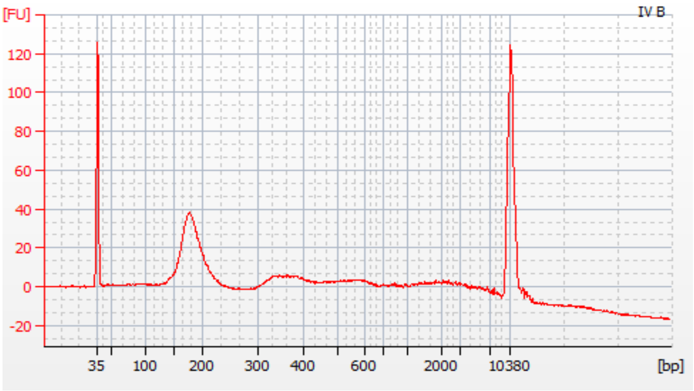  |
| #V | Cell-free DNA | Glioblastoma | 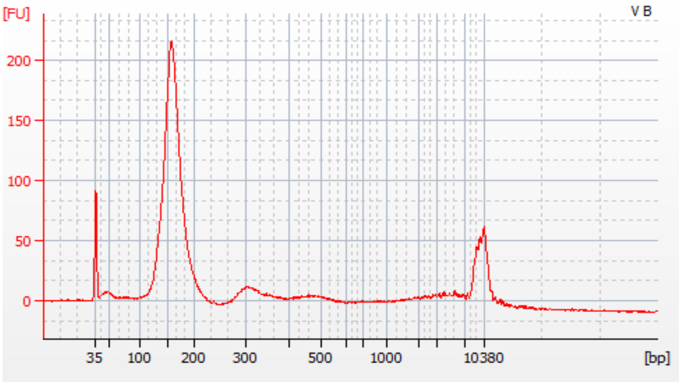  |
| #VIII | Cell-free DNA | Glioblastoma | 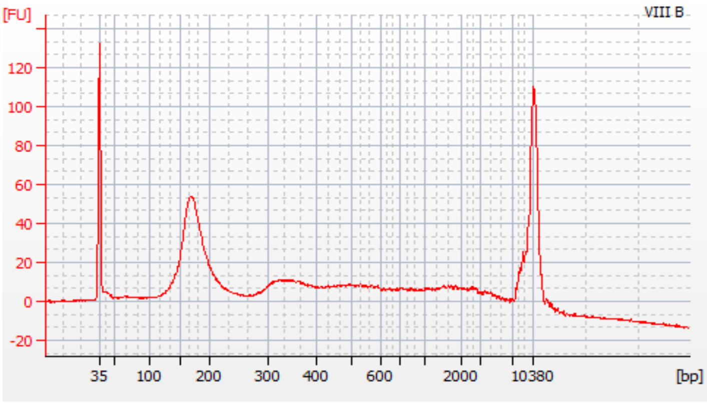  |

| | | | |
|---|---|---|---|
| #IX | Cell-free DNA | Glioblastoma | 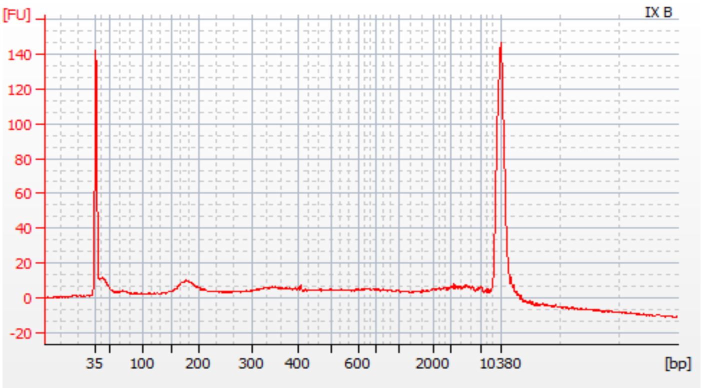 |
| #X | Cell-free DNA | Glioblastoma | 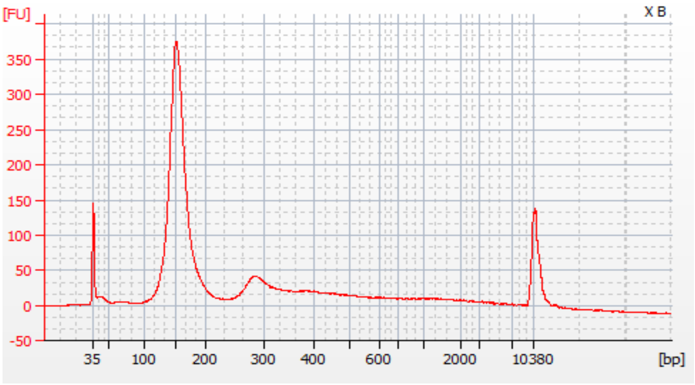 |
| #XI | Cell-free DNA | Glioblastoma | 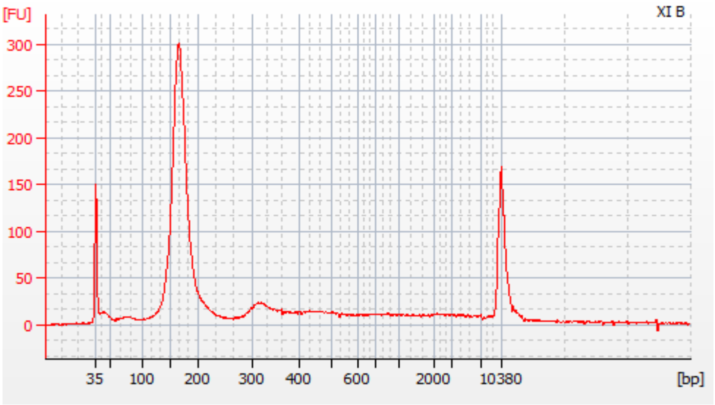 |

| | | | |
|---|---|---|---|
| #XII | Cell-free DNA | Glioblastoma | 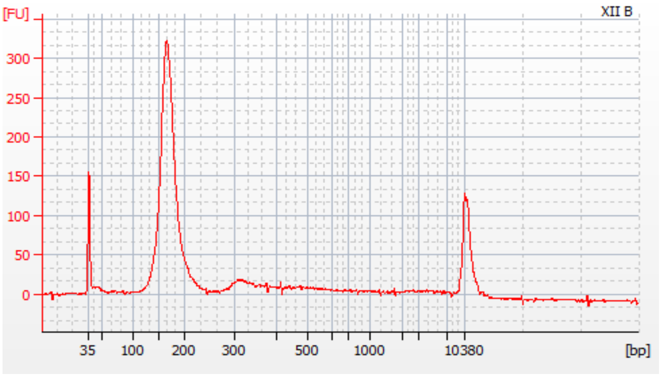 |
| #592 | Cell-free DNA | Low-grade glioma | 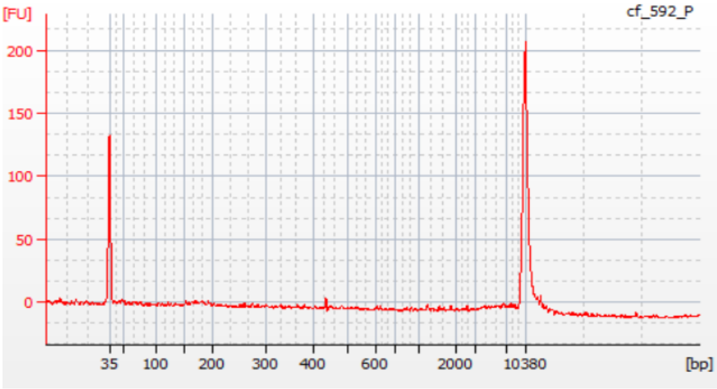 |
| #915 | Cell-free DNA | Low-grade glioma | |

| | | | |
|---|---|---|---|
| #Les-S | Cell-free DNA | Healthy Control | 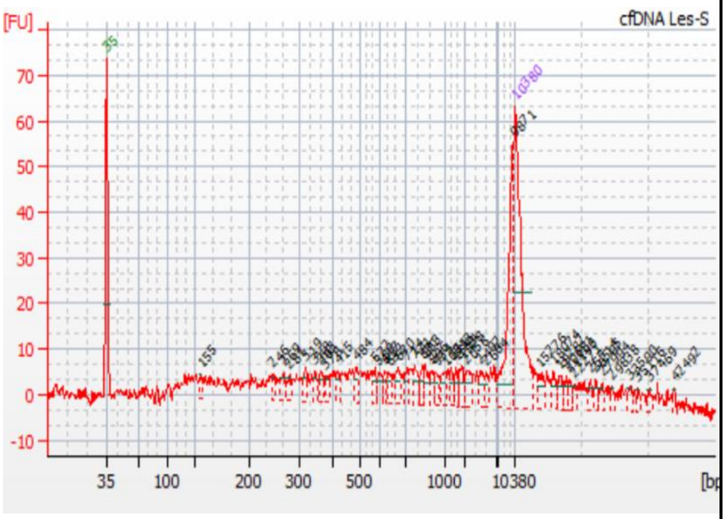 |
| #Les-CM | Cell-free DNA | Healthy Control | 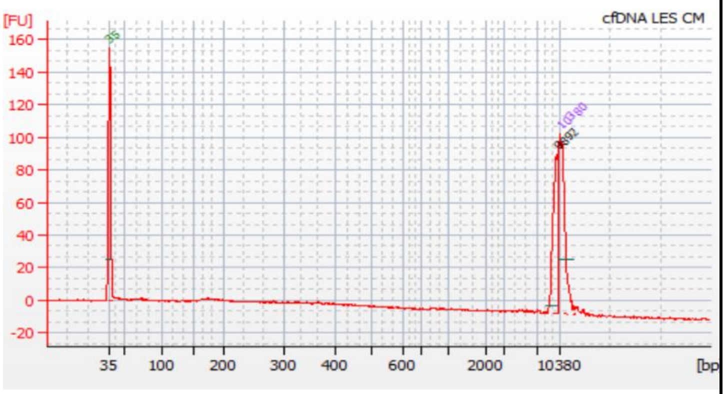 |
| #LL | Cell-free DNA | Healthy Control | 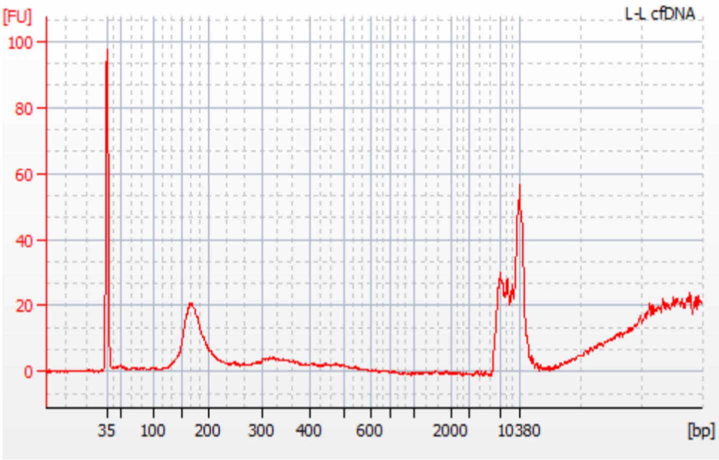 |

| | | | |
|---|---|---|---|
| #CH-L | Cell-free DNA | Healthy Control | 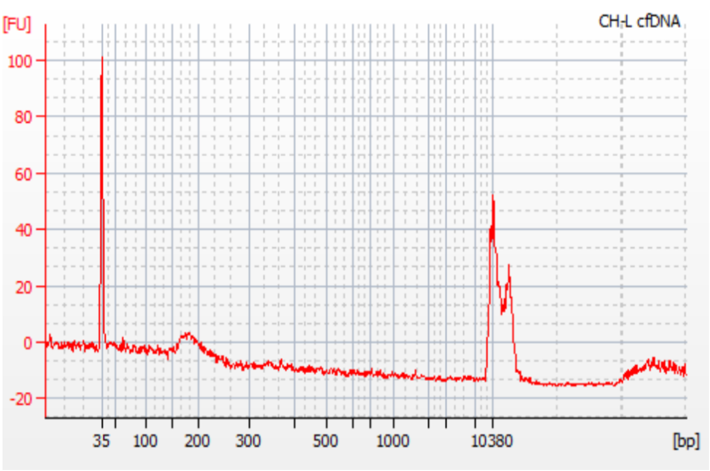 |
| #TS-105 | Cell-free DNA | Healthy Control | 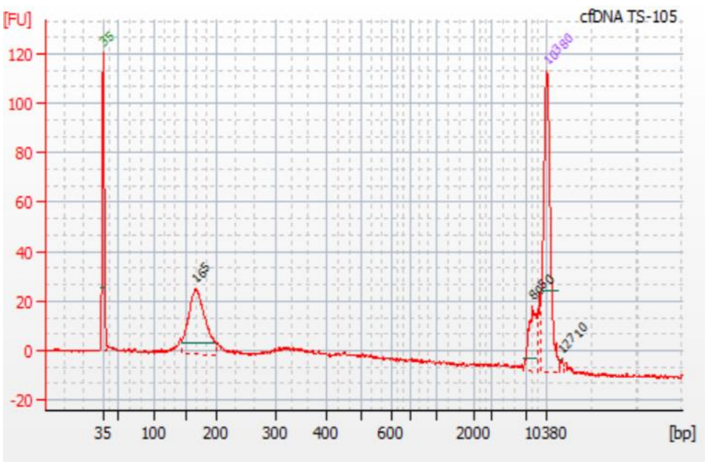 |
| #TS-IN | Cell-free DNA | Healthy Control | 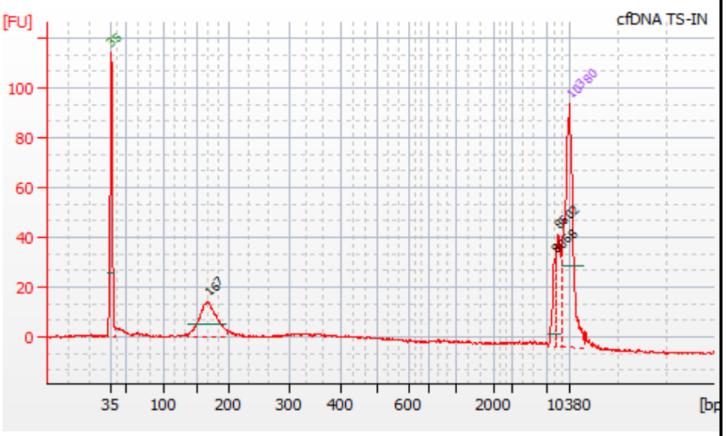 |

| | | | |
|---|---|---|---|
| #TS-LID | Cell-free DNA | Healthy Control | 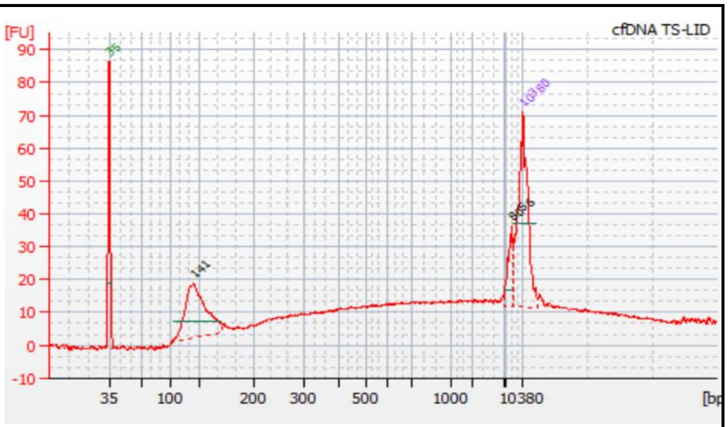 |
| #TS-0 | Cell-free DNA | Healthy Control | 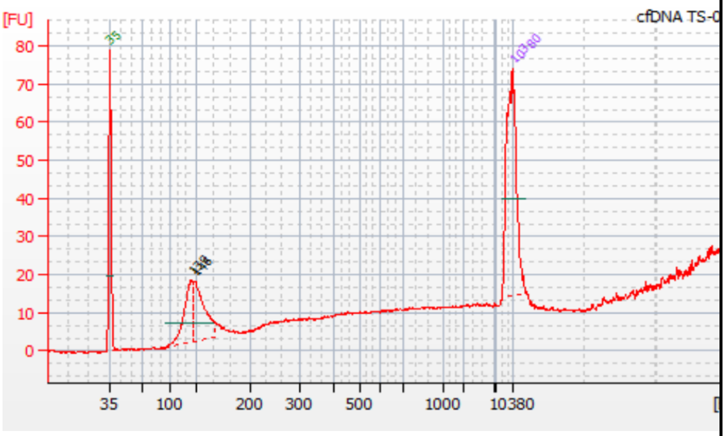 |
| #TS-IL | Cell-free DNA | Healthy Control | 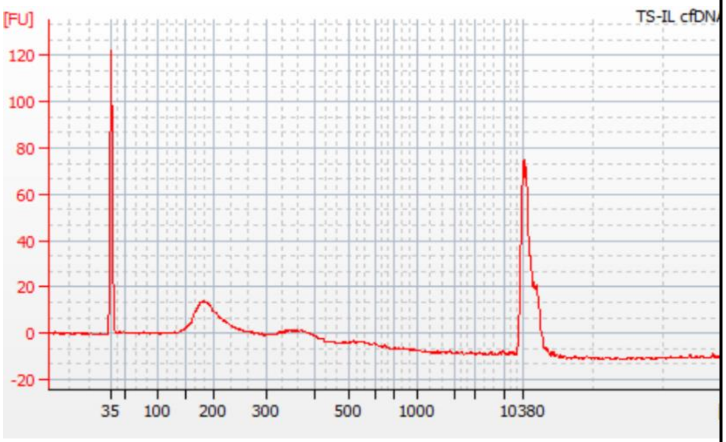 |
